# Apoplastic barrier establishment in roots and nodules of Lotus japonicus is essential for root-shoot signaling and N-fixation

**DOI:** 10.1101/2023.12.06.570432

**Authors:** Defeng Shen, Rafael E. Venado, Ulla Neumann, Nadine Dyballa-Rukes, Swati Mahiwal, Sabine Metzger, Ryohei Thomas Nakano, Macarena Marín, Tonni Grube Andersen

## Abstract

The molecular framework underlying apoplastic root barrier formation has been unveiled in the model species *Arabidopsis thaliana* where establishment of Casparian strips occurs at an early stage of root development. In legumes, this region overlaps with the area where nitrogen-fixing bacteria can induce nodule formation, termed the susceptible zone. Moreover, while nodules themselves also contain an endodermis spanning their vascular bundles, it is current unknown if Casparian strips serve as a filter for transport across this specialized organ. Here we establish barrier mutants in the symbiosis model *Lotus japonicus.* We find that the while genetic network controlling Casparian strip formation is conserved in this legume species, formation of functional barriers is crucial for establishment of N-fixing nodules. By probing this in detail, we establish a model where the Casparian strip, via its linked Schengen pathway, converge with long distance N signaling and systemic regulation of nodulation. Moreover, this also reveal that the genetic system for barrier establishment in the root endodermis is shared in nodule vascular endodermis and required for nodule function. Combined, our findings uncover a novel role of apoplastic root barriers and establishes a mutant collection suitable to probe the role of root barriers in symbiotic plant-microbe relationships.

## Introduction

In vascular plants, the root endodermis provides a mechanism that filters solutes by forcing uptake to occur via transport across the plasma membrane (*1*). This emerges through localized lignin depositions known as Casparian strips (CSs), situated in the cell walls between endodermal cells (*1*). While the CS was discovered almost two centuries ago (*2*), the underlining molecular mechanism responsible for its formation has only recently been unveiled. Based on work in the model species *Arabidopsis thaliana* (Arabidopsis), we now have a detailed understanding of how CS deposition is facilitated in young endodermal cells (*3*). In short, an initial localization of CASPARIAN STRIP DOMAIN PROTEINs (CASPs) at the CS membrane domain (CSD) acts as a scaffold that recruits proteins necessary for lignin biosynthesis, such as ENHANCED SUBERIN1 (ESB1) and a number of PEROXIDASEs (PERs) (*4–6*). The genetic network responsible for CS establishment is controlled by the R3R2 MYB-class transcription factor MYB36 (*7*, *8*). However, to support the formation of a coherent, functional barrier the MYB36-driven CS machinery is dependent on a signaling pathway termed the Schengen (SGN) pathway. This “surveillance” system consists of the stele-produced CASPARIAN STRIP INTEGRITY FACTOR (CIF) peptide ligands, the leucine-rich repeat receptor-like kinase (LRR-RLK) SCHENGEN3/GASSHO1 (SGN3/GSO1) and the downstream receptor-like kinase SCHENGEN1 (SGN1/PBL15) (*9–11*). During CS establishment, CIF peptides diffuse to the cortex-facing side of the endodermis where they bind a complex consisting of SGN3 and SGN1. This activates downstream pathways, which ultimately lead to CS fusion and thereby inhibition of further SGN activation (*9*, *11*). In CS defective mutants (e.g. *myb36* knockouts), increased diffusion of CIF peptides results in a hyperactive SGN response, which onsets ectopic endodermal suberization and lignification (*5*, *7*). Evidence is emerging that orthologous genes involved in CS formation play a similar role in other plants (*12–17*). We recently identified the Arabidopsis SGN pathway to play a crucial role in integrating the soil nitrogen (N) status into shoot responses (*18*). Combined, this raises the question whether the CS-SGN coordination of environmental signals is present in other species.

N-related long-distance signaling between roots and shoots are especially important in plants that form a symbiotic relationship with N-fixing bacteria through establishment of specialized root-organs termed nodules (*19*). In model species such as *Medicago truncatula* and *Lotus japonicus* (hereafter Lotus), nodule formation is tightly regulated by N availability and controlled by a systemic signaling system called the autoregulation of nodulation (AON) (*19*). Under low N conditions, C-TERMINALLY ENCODED PEPTIDE1 (CEP1) peptides are induced in the root stele (*20*), translocated to the shoot and perceived by the COMPACT ROOT ARCHITECTURE2 (CRA2) receptor (*21*). Activated CRA2 receptors in the shoots induce the production of phloem-mobile signals known as CEP DOWNSTREAM 1 and 2 (CEPD1 and 2) (*22*). Besides this, CRA2 also promotes the synthesis of microRNA2111 (miR2111), which moves to the roots and inhibits translation of the nodulation repressor *TOO MUCH LOVE* (*TML*) (*22*, *23*). Combined, this allows nodules to form specifically when the plant cannot obtain sufficient N from the surrounding soil.

Initiation of nodule formation is restricted to a narrow developmental window of the root termed the susceptible zone (SZ)(*24*). Intriguingly, this zone is situated closer to the root tip, overlaps with the region in which the functional CS is established (*4*). As nodule is primarily formed by root cortex (*25*), this implies that communications via the vasculature for the systemic control of nodulation occur through root endodermis, and a likely coordination with CS formation. Moreover, within the mature nodule, the nodule vasculature is surrounded by endodermal cells that contain CS (*26*). Thus, exchange of bacterially fixed N and photosynthates between symbionts and the nodule vasculature must be subject to apoplastic regulation analogously to the scenario of endodermal filtering in roots.

Here we address these questions by identifying mutants with defective root barrier formation in the legume model Lotus. We employ these to study the CS-SGN system in connection to N-related signaling, nodule formation and function. Combined, our work demonstrates an important function of the CS-SGN system in systemic signaling underlying N-related symbiosis. This not only confirms an essential role of the CS-SGN system in long-distance N-signaling, but establishes apoplastic barriers as an important mechanism in spatially constrained symbiotic plant-microbe relationships.

## Results

### CS establishment network is conserved in Lotus

To identify orthologues of verified CS-related genes from Arabidopsis in Lotus, we performed a phylogenetic analysis on *MYB36*, SGN1 and *SGN3*. This identified the gene LotjaGi4g1v0298100 as the putative orthologue of *AtMYB36* (hereafter *LjMYB36*) and LotjaGi2g1v0345700 as the putative orthologue of *AtSGN1* (hereafter *LjSGN1*). For SGN3 we found two putative orthologues: LotjaGi4g1v0018600 (hereafter *LjSGN3a*) and LotjaGi6g1v0246100 (hereafter *LjSGN3b*) (Supplementary Figure 1A) (Supplementary Table 1). To investigate if these genes play similar roles for CS formation as in Arabidopsis, we obtained *LOTUS RETROTRANSPOSON 1* (*LORE1*) knockout (KO) mutant alleles for *LjMYB36*, *LjSGN1*, *LjSGN3a* and *LjSGN3b* (*27*) (Supplementary Table 2). We successfully isolated two independent homozygous lines for each of these genes, except for *LjSGN3b.* In Arabidopsis, the highly similar homolog of AtSGN3, AtGSO2 is involved in embryo development (*28*). Therefore, LjSGN3b might be the orthologue of AtGSO2 and possibly serves an essential function in Lotus, rendering homozygous mutants non-viable. In support of a CS-related function, the promoter regions of *LjMYB36*, *LjSGN1* and *LjSGN3a* showed specific expression in the differentiation zone concurrent with the establishment of CS (Supplementary Figure 1B). Next, we investigated if CS formation was affected in the remaining mutants. A recent study reported a *Ljmyb36-1* mutant (*29*), we therefore termed our *Ljmyb36* mutants *Ljmyb36-2* and *Ljmyb36-3*. Gifu (background of *LORE1* mutants) and all segregated wild-type (WT) lines showed lignin-specific staining consistent with CS formation, this was absent in all KO mutant lines (Figure 1A, Supplementary Figure 1C). Moreover, in support of disturbed CS function, these mutants had a strong delay in blockage of the apoplastic tracer propidium iodide (PI) (*30*) (Figure 1B and C, Supplementary Figure 1D). As *Ljsgn3a* clearly demonstrated a CS defective phenotype, we hereafter termed *LjSGN3a* as *LjSGN3*. Transgenic roots expressing genomic sequence of *LjMYB36* or coding sequence of *LjSGN1* and *LjSGN3* using native promoter and 3’ untranslated region can complement CS formation in the respective mutant background (Supplementary Figure 1E). Combined, we conclude that *LjMYB36*, *LjSGN1* and *LjSGN3* are all essential for functional CS formation in Lotus roots.

**Figure 1.**
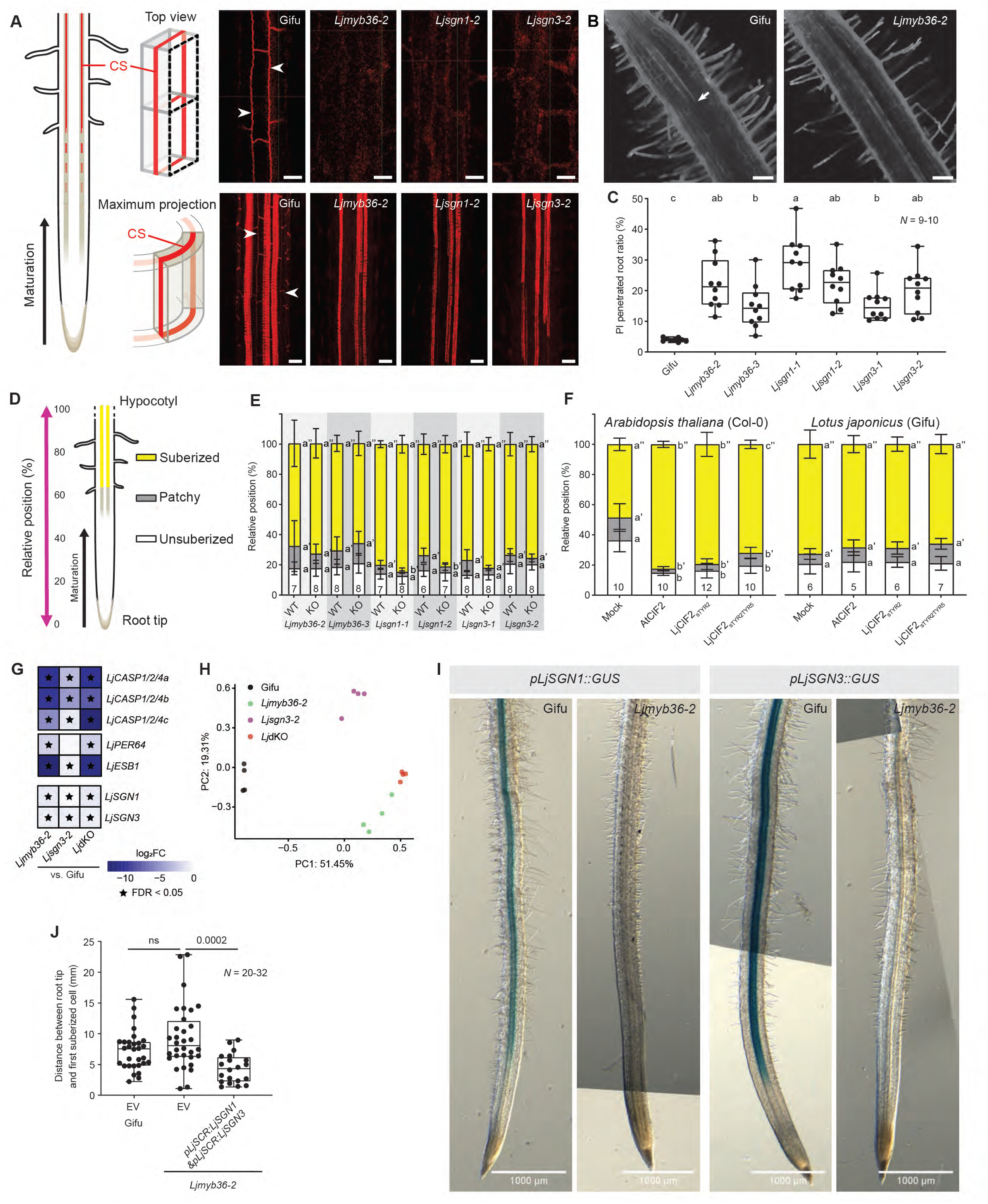
Characterization of Lotus Casparian strip defective mutants. (**A**) Top view (upper) and Maximum projection (lower) of confocal image stacks of Basic Fuchsin stained 9-day-old Gifu, *Ljmyb36-2*, *Ljsgn1-2* and *Ljsgn3-2* roots at a similar region. Arrowhead indicates Casparian strip (CS). Representative images from three independent experiments (n=18). (**B**) Representative images of propidium Iodide (PI) stained 9-day-old Gifu and *Ljmyb36-2* roots. Arrow indicates the blockage of PI penetration (arrow) into the vascular bundle. (**C**) Quantification of the proportion of 9-day-old roots can be penetrated by PI into the vascular bundle. Representative results from two independent experiments. (**D**) Representation of suberization pattern in Lotus roots, with unsuberized zone closer to the root tip, followed by patchy zone and continuous suberized zone in the direction of hypocotyl. See Supplementary Figure 1F for an example. (**E**) Suberization pattern of 8-day-old Lotus CS defective mutants, compared with their corresponding segregated wild-type (WT) lines. Different letters depict the statistical difference between mutant with their corresponding segregated WT lines in a two-sided Student’s t test (*P* < 0.05). Representative results from two independent experiments. (**F**) Left: suberization pattern of 7-day-old Arabidopsis Col-0 roots after 48 h treatment with 100 nM CIF2 peptide under agar condition. Right: suberization pattern of 8-day-old Lotus Gifu roots after 48 h treatment with 500 nM CIF2 peptide under agar condition. Representative results from two independent experiments. (**G**) Heatmap of genes involved in CS formation in Lotus CS defective mutants. (**H**) Principal coordinate analysis (PCoA) plot of transcriptome of Lotus CS defective mutants. (**I**) Transcriptional activity of *LjSGN1* (left) and *LjSGN3* (right) in Gifu and *Ljmyb36-2* hairy roots, visualized by GUS signal. Representative images of at least 10 composite plants. (**J**) Quantification of distance between root tip and first suberized cell in *Ljmyb36-2* hairy roots overexpressing *LjSGN1* and *LjSGN3* using *pLjSCR*, compared with transgenic controls (EV, *pLjSCR:GUS*) of Gifu and *Ljmyb36-2*. Combined results of two independent experiments. *P* values indicate two-sided Student’s t test (ns, not significant). Numbers of plants investigated are indicated in graph. Different letters in (**C**) (**F**) depict the statistical difference in a one-way ANOVA analysis with Tukey’s test (*P* < 0.05), unless otherwise indicated. Scale bar: 20 µm (**A**), 100 µm (**B**).

### Different suberization pattern in Lotus CS defective mutants

In Arabidopsis, disruption of CS formation leads to SGN-dependent ectopic lignification and early onset of endodermal suberization (*3*). Remarkably, these compensatory effects were not observed in either of the two *Ljmyb36* mutant alleles (Figure 1D and 1E, Supplementary Figure 1F). As this indicates differences in the SGN pathway, we questioned whether the Lotus SGN signaling can be activated by exogenous CIF peptides. We identified two Lotus genes encoding putative CIF peptides (Supplementary Figure 2A) and investigated their expression in publicly available transcriptome datasets (Lotus Base) (*31*). Similar to Arabidopsis (*10*), one of the Lotus CIF peptide-encoding genes had a higher expression level (approximately 20-fold) than the other in roots (Supplementary Figure 2B). We therefore named the higher expressed CIF peptide encoding gene as *LjCIF2*, the other one as *LjCIF1*. In Arabidopsis, external application of CIF peptide leads to increased lignin and suberin deposition (*9*). Both mature LjCIF peptides contain two putatively sulfated tyrosine residues (Tyr) at position 2 and 5, compared with only one in AtCIF at position 2 (Supplementary Figure 2A). To test if CIF peptides can induce ectopic SGN responses in Lotus, and account for the different sulfation pattern, we synthesized LjCIF2_sTyr2_ (sulfation at Tyr2) and LjCIF2_sTyr2Tyr5_ (sulfations at Tyr2 and Tyr5). When applied to Arabidopsis both peptides were able to induce overlignification and oversuberization at similar concentrations as AtCIF2 (Figure 1F, Supplementary Figure 2C), demonstrating them as *bona fide* ligands for AtSGN3. However, none of the CIF peptides could induce ectopic suberization or lignification in Lotus Gifu roots, even when applied at higher concentrations (Figure 1F, Supplementary Figure 2C). As Lotus has thicker roots than Arabidopsis, we checked if the extra cell layers may prevent external CIF diffusion from reaching the endodermis. This does not appear to be the case as we observed similar lack of responses when adding peptides to plants grown in a hydroponic growth setup (Supplementary Figure 2D) and fluorescently labeled AtCIF2 (TAMRA-AtCIF2) peptide was able to reach the endodermis in the differentiated root (Supplementary Figure 2E). Therefore, we conclude that in Lotus the SGN pathway is involved in CS formation, but cannot be activated by exogenously applied CIF ligands.

### Transcriptomic analysis of Lotus Casparian strip mutant roots

To further investigate the differences in SGN function, we performed a transcriptional analysis of these CS defective mutants. As the *Ljmyb36* KO lines phenocopied the *Atmyb36sgn3* double knockout (dKO) with regards to the diminished SGN response, we created a *Ljmyb36-2xsgn3-2* dKO by crossing these two lines (hereafter *Lj*dKO). This allowed us to directly compare the hierarchy of the underlying transcriptional networks in Lotus with a similar dataset in Arabidopsis (*32*). Indeed, similarly to Arabidopsis, genes involved in CS formation (e.g. *LjCASPs*, *LjESB1* and *LjPER64*) were repressed in *Ljmyb36-2*, *Ljsgn3-2* as well as the *Lj*dKO roots, compared to Gifu roots (Figure 1G, Supplementary Figures 3A). This is consistent with the observed defective barrier formation in those mutants (for *Lj*dKO, see Supplementary Figures 1C, 1D and 3B). By performing a Gene Ontology (GO) term analysis, we found that none of the SGN responses which has been described to occur in *Atmyb36-2* (e.g. phenylpropanoid biosynthetic process) (*32*) was activated in *Ljmyb36-2* roots (Supplementary Figure 3C). A principal coordinate analysis (PCoA) revealed that in contrast to Arabidopsis, the *Lj*dKO clustered with the *Ljmyb36-2* (Figure 1H, Supplementary Figure 3D), which shows that little additional transcriptional variation occurred by knocking out *LjSGN3* in *Ljmyb36-2* background. One interpretation of this is that the SGN pathway may depend on MYB36 in Lotus. Indeed, the expression of both *LjSGN1* and *LjSGN3* was strongly repressed in *Ljmyb36-2* roots when compared to Gifu, and virtually absent when using β-glucuronidase (GUS)-based reporters (Figure 1I). Moreover, overexpression of both of *LjSGN1* and *LjSGN3* by the endodermis-specific Lotus *SCARECROW* (*LjSCR*) promoter (*33*) induced earlier suberization in *Ljmyb36-2* roots when compared to control roots (Figure 1J). Combined, we therefore conclude that in Lotus the SGN pathway is dependent on MYB36 and the lack of SGN response in *Ljmyb36* roots is due to the attenuated expression of *SGN* genes.

### Arabidopsis and Lotus SGN3 display differences in CIF recognition

To evaluate how conserved the functionality of the SGN1/3 module is between Arabidopsis and Lotus, we complemented the corresponding Arabidopsis mutants with their native or Lotus versions of SGN1 or SGN3. Both AtSGN1 and LjSGN1 were able to fully complement the dysfunctional CS as well as suberin induction by CIF2 peptides in the *Atsgn1*-2 mutant (Figure 2A, Supplementary Figure 4A). For SGN3, however, only AtSGN3 could complement CS formation in the *Atsgn3-3* mutant, whereas lines expressing LjSGN3 still contained a dysfunctional barrier (Figure 2A). Intriguingly, while externally applied AtCIF2 and LjCIF2_sTyr2_ were able to induce ectopic suberization in all complementation lines, peptides induced a stronger change in *Atsgn3-3* lines complemented with the native SGN3 receptor (Figure 2B). Thus, LjSGN3 appears to be functional in the complemented lines, but display selectivity towards its endogenous CIF peptide in relation to induction of ectopic suberin. Like most LRR-type receptor-like kinases, SGN3 consists of a ligand-binding ectodomain as well as an intracellular kinase domain that facilitates downstream signaling via phosphorylation (*34*). Thus, differences in CIF recognition likely lie in the LRR ectodomain. In support of this, *Atsgn3-3* lines complemented with a chimera of AtSGN3 (ectodomain) and LjSGN3 (kinase domain) could fully complement the defective CS formation in *Atsgn3-3* (Supplementary Figure 4B), which proposes that the perception and binding of CIF peptides in the ectodomain is different between these two species.

**Figure 2.**
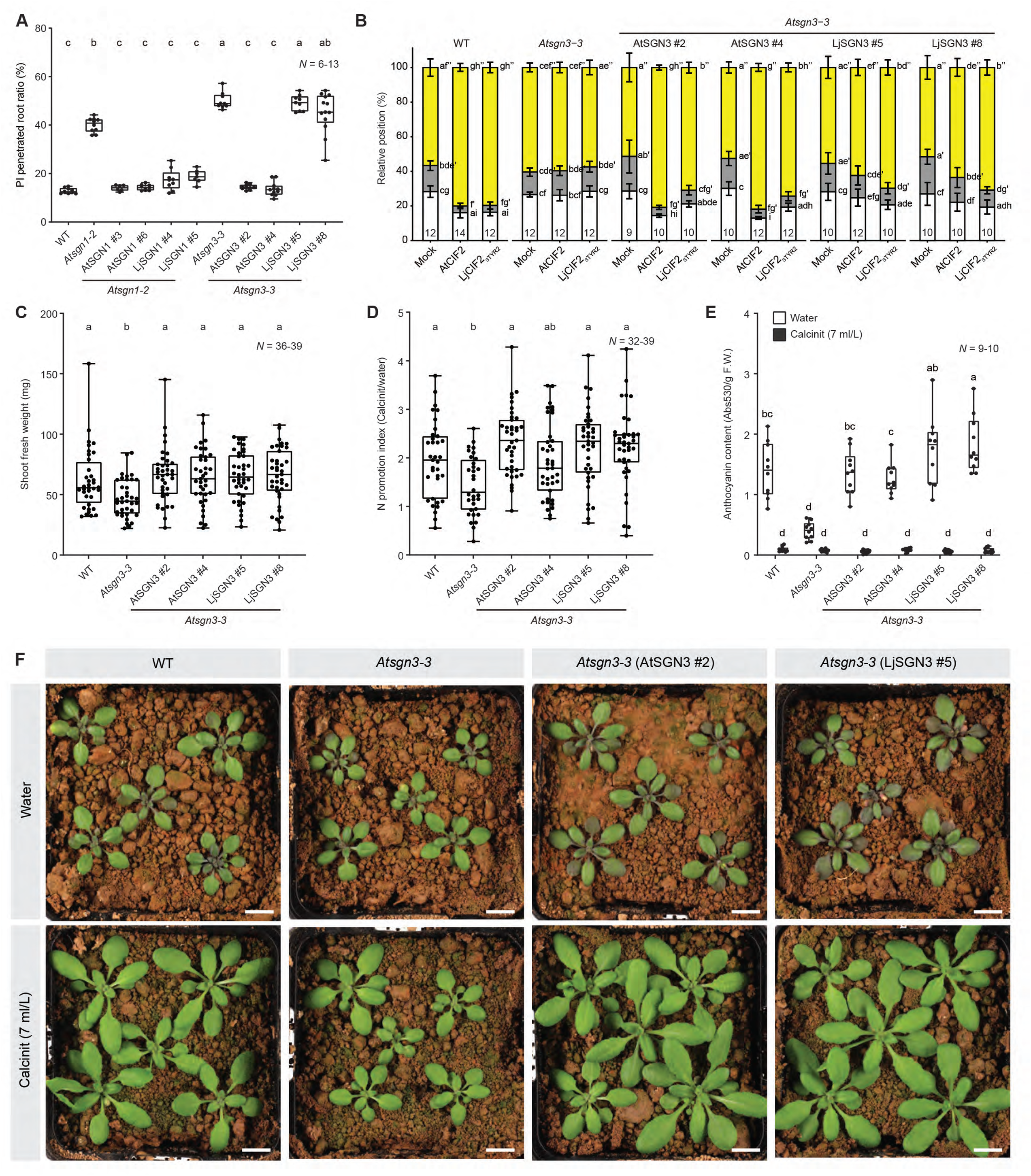
Conserved function shared by SGN3 in Arabidopsis and Lotus. (**A**) Quantification of PI penetration in 7-day-old roots of *Atsgn1-2* and *Atsgn3-3* complementation lines. (**B**) Suberization pattern in 7-day-old roots of *Atsgn3-3* complementation lines treated with mock, AtCIF2, or LjCIF2_sTYR2_ for 48 h. (**C**) Shoot fresh weight under water condition, (**D**) Nitrogen promotion index (Calcinit treated shoot fresh weight/water treated shoot fresh weight), (**E**) anthocyanin content, (**F)** representative images of 28-day-old *Atsgn3-3* complementation lines grown in CAS. Different letters depict the statistical difference in a one-way ANOVA analysis with Tukey’s test (*P* < 0.05). (**A**-**F**) Representative results from two independent experiments. Scale bar: 1 cm (**F**).

### SGN3-dependent N-induced growth promotion is independent of Casparian strip status

Recently, we found evidence in Arabidopsis that SGN3 plays a role in N-induced growth promotion under agricultural soil conditions (*18*). To test if this effect is conserved, we evaluated growth of AtSGN3- and LjSGN3-complemented *Atsgn3-3* lines under a N-fertilization scheme using our local Cologne agricultural soil (CAS) (*35*, *36*). As previously described (*18*), when compared to WT, *Atsgn3-3* displayed a diminished N-induced growth promotion, decreased growth under low N conditions and reduced N-starvation-induced anthocyanin accumulation in the shoots when compared to WT (Figure 2C-F). Remarkably, both *AtSGN3* and *LjSGN3*-expressing lines fully complemented these responses despite the lack of functional CS formation in the LjSGN3-complemented lines (Figure 2C-F). Thus, while the CS establishment machinery is distinct, the ability of SGN3 to activate shoot responses related to soil N status is conserved between Arabidopsis and Lotus.

### Casparian strip establishment in the root facilitates nodule formation

Since the root region where CS is established overlaps with the SZ of rhizobia, we questioned whether nodule formation is affected in Lotus CS defective mutants. Indeed, nodule numbers were significantly reduced on *Ljmyb36, Ljsgn1* and *Ljsgn3* mutant roots when compared to Gifu and segregated WT lines (Figure 3A and B, Supplementary Figure 5A). Moreover, In Gifu the first nodule on the primary root occurred immediately above or at the root tip position marked at 0 day post-inoculation (dpi) (Figure 3A and C). In contrast, all CS mutants showed a delayed nodule formation accompanied by a significant reduction in nodule size (Figure 3A and C, Supplementary Figure 5B). Moreover, the infection thread density was significantly reduced in CS mutants at 5 dpi and 9 dpi (Supplementary Figure 5C). In addition to this, while spot inoculation of rhizobia induced cell divisions on mutant roots at 3 dpi, only a minority formed nodules at 14 dpi (Figure 3D-F). Thus, besides negatively influencing the infection procedure, the barrier mutants were additionally defective in mechanisms that facilitate progression of nodule formation.

**Figure 3.**
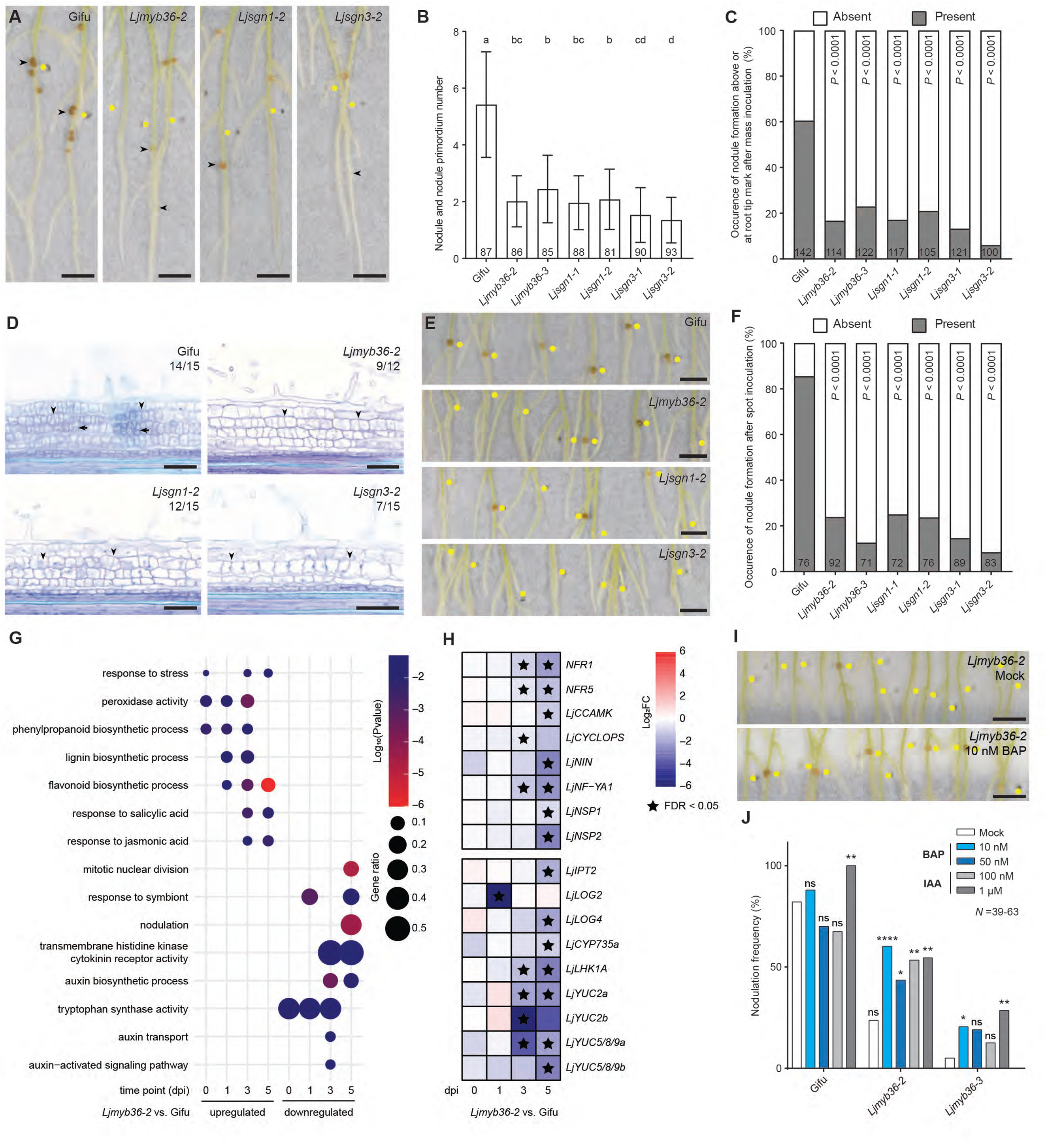
Nodule formation is disturbed in CS defective mutant roots. **(A)** Representative images of mass-inoculated roots with nodules at 21-days post-inoculation (dpi). Arrowheads indicate the first nodule formed on the primary root. (**B**) Quantification of nodule and nodule primordium number at 21 dpi after mass inoculation. Combined results from three independent experiments. Different letters in depict the statistical difference in a one-way ANOVA analysis with Tukey’s test (*P* < 0.05). (**C**) Quantification of occurrence of nodule formed above or at the root tip position marked at 0 DAI after mass inoculation. Combined results from four independent experiments. *P* values indicate Fisher’s exact test (comparison with Gifu). (**D**) Representative images of semi-thin sections on nodule primordia at 3 dpi after spot inoculation. Numbers on the graph indicate proportion of root segments with shown mitotic activity. Arrowheads and arrows indicate anticlinal and periclinal divisions induced in cortex, respectively. (**E**) Representative images of spot-inoculated roots with nodules at 14 dpi. (**F**) Quantification of occurrence of nodule formation at 14 dpi after spot inoculation. Combined results from three independent experiments. *P* values indicate Fisher’s exact test (comparison with Gifu). (**G**) Dotplot of Gene Ontology terms enriched in upregulated and downregulated DEGs of *Ljmyb36-2* nodule primordia at different time points after spot inoculation. (**H**) Heatmap of symbiosis related genes (top), auxin and cytokinin related genes (bottom, for full list see Supplementary Figure 6E) in *Ljmyb36-2* nodule primordia at different time points. (**I**) Representative images of spot-inoculated *Ljmyb36-2* roots supplemented with mock (top) or 10 nM BAP (bottom) at 14 dpi. (**J**) Quantification of occurrence of nodule formation at 14 dpi after spot inoculation. Fisher’s exact test was performed between hormone treatment and mock within the same genotype (ns, not significant, * *P* < 0.05, ** *P* < 0.01, **** *P* < 0.0001). Combined results from two independent experiments. Yellow dots indicate the position of primary root tip at 0 dpi. Numbers on the graph indicate the number of plants investigated. Scale bar: 5 mm (**A, E, I**), 100 µm (**D**). dpi: day post-inoculation.

### Disturbed hormone signaling homeostasis in Casparian strip mutant nodule primordia

To explore the molecular basis of aborted nodule formation in the Lotus CS mutants, we performed a time-resolved whole-transcriptome analysis of spot-inoculated Gifu, *Ljmyb36-2* and *Ljsgn1-2* roots at 0, 1, 3, 5 dpi. A PCoA on all samples revealed that the developmental stage of nodule primordia has a stronger effect on the overall root transcriptome rather than the presence of functional CS (Supplementary Figure 6A). However, when we divided the samples into individual time points, we observed distinct transcriptional responses between Gifu and CS defective mutants (Supplementary Figure 6B). By performing a Gene Ontology (GO) term analysis, we found that “response to symbiont” and “nodulation” were enriched in the downregulated DEGs (FDR-adjusted *P* value < 0.05, absolute fold change >2) found in the CS defective mutants (Figure 3G, Supplementary Figure 6C). In accordance, genes that positively regulate nodulation (e.g. *NFR1, NFR5, CCAMK, NIN, NF−YA1*) were downregulated in *Ljmyb36-2* and *Ljsgn1-2* nodule primordia at 3 dpi and/or 5 dpi (Figure 3H, Supplementary Figure 6D). GO terms related to “mitotic nuclear division” as well as auxin and cytokinin homeostasis were overrepresented in the reduced DEGs in *Ljmyb36-2* and *Ljsgn1-2* nodule primordia (Figure 3G and H, Supplementary Figure 6C and E). This corroborated our histological analysis and suggested that one of the underlying causes for the suppressed nodule formation is disturbed hormonal balances. Indeed, mutant roots spot-inoculated with rhizobia and supplemented with either the artificial cytokinin 6-benzylaminopurine (BAP) (10 nM and 50 nM) or the auxin Indole-3-actic acid (IAA) (1 µM and 10 µM) showed a significant increase in nodulation frequency in *Ljmyb36*, but not in *Ljsgn1* or *Ljsgn3* mutants (Figure 3I and J, Supplementary Figure 6F). Thus, the aborted cell division in *Ljmyb36* nodule primordia is likely related to local changes in cytokinin and auxin homeostasis, which may depend on the SGN signaling system.

### Root-shoot communication for nodule formation is dysfunctional in Lotus CS mutants

Next, we sought a common, upstream explanation for the reduced nodulation among our CS mutants. Our transcriptome analysis revealed that the central player of AON, *TML* (*19*) was upregulated in *Ljmyb36-2*, *Ljsgn3-2* and *Lj*dKO roots (Supplementary Figure 3E) as well as in the SZ of *Ljmyb36-2* roots at 0 dpi (Supplementary Figure 6G). Thus, these mutants may have dysfunctional AON regulation and disturbed systemic N-signaling. Indeed, homozygous crossings of the hyper-nodulating *tml5* KO mutant (*23*) with *Ljmyb36-2* and *Ljsgn3-2* markedly promoted nodulation and induced earlier nodulation events (Figure 4A-C, Supplementary Figure 7A). Next, we investigated how *TML* expression was mis-regulated in CS mutant roots. Since CEP1 peptides influence *TML*, we applied exogenous LjCEP1 peptides to mutant roots, which significantly increased nodule occurrence and induced earlier nodulation across CS defective mutants (Figure 4D-F, Supplementary Figure 7B). As we observed similar results when LjCEP1 was directly applied to mutant shoots (Supplementary Figure 7C), we conclude that these peptides can move to shoots and induce downward signals to promote nodulation despite the dysfunctional CS in the mutants. We further found that under axenic conditions miR2111 abundance was significantly reduced in *Ljmyb36-2* shoots and SZ, while *TML* transcripts were increased accordingly in *Ljmyb36-2* SZ (Figure 4G-I). This could be explained by reduced expression of *LjCEP1* in *Ljmyb36-2* roots (Figure 4J). In further support of this, the expression of *LjCEPD1* (activated by LjCEP1 in shoots) (*22*) was also reduced in *Ljmyb36-2* shoots (Figure 4K). This indicates that the disturbed CS affects expression of the gene encoding for the signaling peptide rather than affecting the upward movement, as may be expected from a mutant with defective CS. We further found that application of exogenous LjCEP1 to the roots led to significantly increased expression of *LjCEPD1* in both Gifu and *Ljmyb36-2* shoots (Figure 4K). This also led to a higher abundance of miR2111 and reduced expression of *TML* in *Ljmyb36-2* SZ (Figure 4H and I). To further substantiate this, we ectopically expressed *LjCEP1* using the meristematic stele-specific Lotus *SHORT-ROOT* (*LjSHR*) promoter (*33*), and this gave rise to a significant increase of nodule number in *Ljmyb36-2* roots (Figure 4L and M, Supplementary Figure 7D). Combined, we conclude that disturbed establishment of a functional CS-SGN system affects the ability of the plant to convey long-distance N signaling by reduced expression of *CEP1* (Figure 4N).

**Figure 4.**
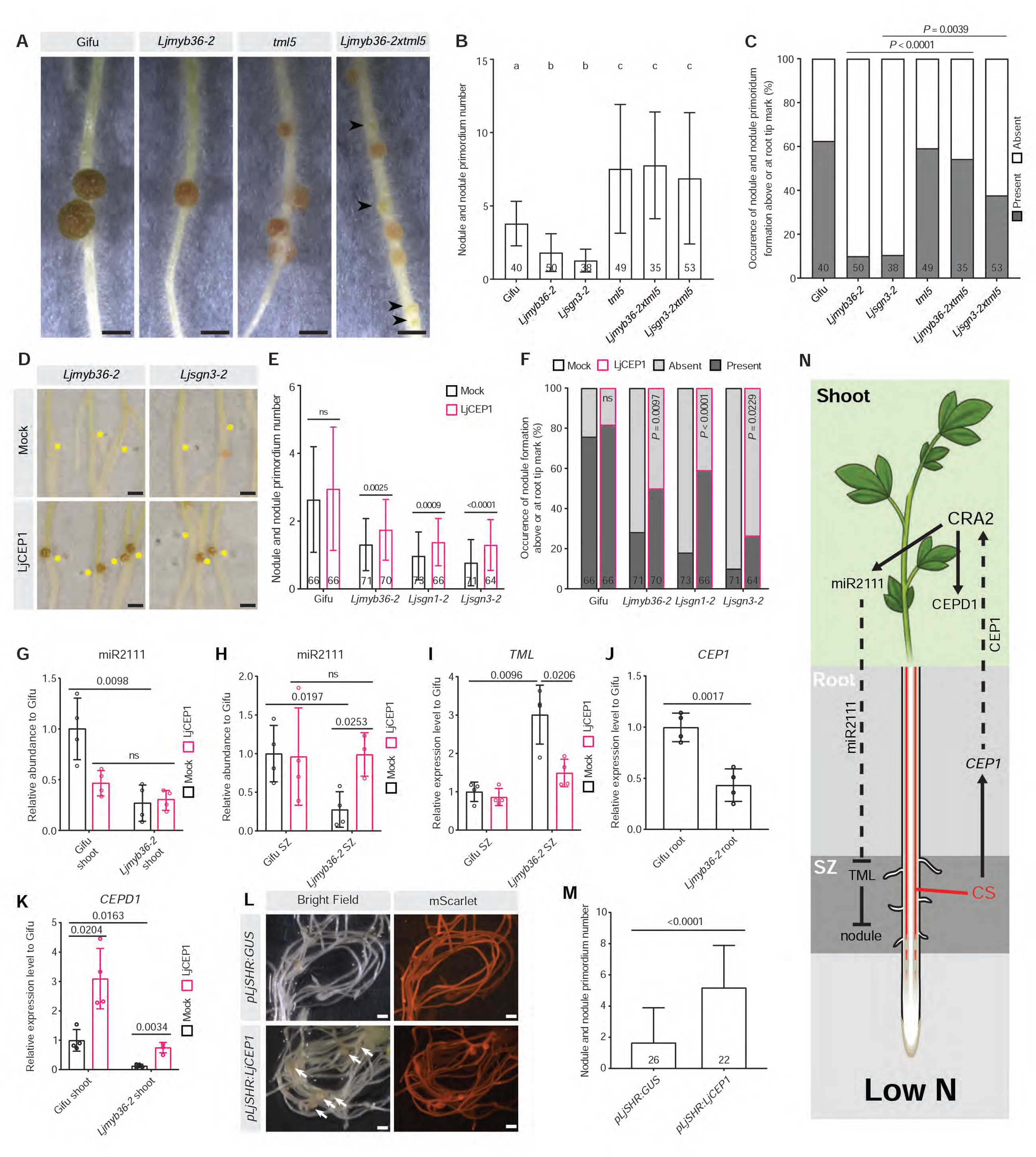
Disturbed nitrogen signaling leads to reduced nodule formation in Casparian strip defective mutant roots. **(A)** Representative images of mass-inoculated roots with nodules at 28 days post-inoculation (dpi). Arrowheads indicate nodule primordia. (**B**) Quantification of nodule and nodule primordium number at 28 dpi after mass inoculation. Combined results from two independent experiments. (**C**) Quantification of occurrence of nodule and nodule primordium formed above or at the root tip position marked at 0 dpi after mass inoculation. Combined results from two independent experiments. *P* values indicate Fisher’s exact test. (**D**) Representative images of mass-inoculated roots treated with mock (water) or 1 µM LjCEP1 at 21 dpi. Yellow dots indicate the position of primary root tip at 0 dpi. (**E**) Quantification of nodule and nodule primordium number at 21 dpi after mass inoculation. Combined results from three independent experiments. (**F**) Quantification of occurrence of nodule and nodule primordium formed above or at the root tip position marked at 0 DAI after mass inoculation. *P* values indicate Fisher’s exact test (comparison between LjCEP1 and mock within the same genotype). (**G-K**) Relative expression level of miR2111 (**G**, **H**), *TML* (**I**) and *LjCEP1* (**J**) and *LjCEPD1* (**K**) by qRT-PCR. Representative results from two independent experiments. (**L**) Representative images of *Ljmyb36-2* hairy roots expressing *pLjSHR:GUS* (top) or *pLjSHR:LjCEP1* (bottom) at 21 dpi under bright field (left) and mScarlet channel (right). Arrows indicate nodules. Representative results from two independent experiments. (**M**) Quantification of nodule and nodule primordium number in *Ljmyb36-2* hairy roots expressing *pLjSHR:GUS* or *pLjSHR:LjCEP1*. Numbers on the graph indicate the number of plants investigated. *P* values indicate two-sided Student’s t test, unless otherwise indicated. (**N**) Under low N conditions, CS in the root susceptible zone (SZ) is required to induce N starvation signal (CEP1) expression to activate CRA2 in the shoots. Activated CRA2 leads to expression of *CEPD1* and synthesis of miR2111. The latter moves to the roots, and represses TML in the SZ, priming the uninfected roots for nodule formation. Scale bar: 1 mm (**A, L**), 2 mm (**E**).

### A conserved machinery for Casparian strip formation in the nodule vascular endodermis

The CS mutants are still capable of producing nodules albeit, these occurred at a lower frequency and appeared paler than Gifu (Figure 5A, Supplementary Figure 8A). This proposes an important function for the establishment of CS on nodule function. By using promoter-GUS reporter lines, we found that *LjMYB36*, *LjSGN1* and *LjSGN3* were transcriptionally active at the peripheral and basal regions of young nodules, but restricted to the nodule vascular endodermis (NVE) in the mature nodules (Supplementary Figure 8B-J). Combined, this suggests a common function in nodules and implies that these mutants have dysfunctional apoplastic barriers in the NVE in a similar manner as in the roots. Indeed, transmission electron microscopy of mature nodules (21 dpi) revealed that CS was formed in the NVE of Gifu, but was not detectable in *Ljmyb36-2* nodules (Figure 5B, Supplementary Figure 9E and F). In *Ljsgn1-2* and *Ljsgn3-2* nodules, CS could still be formed sporadically between most of the NVE cells (Supplementary Figure 9A, B, G, and H). To test if a functional barrier forms inside the nodules, we incubated nodules in a solution of PI to assess the blockage of penetration from the outside into the nodule vasculature. While Gifu nodules showed a clear blockage of PI penetration, evident by the lack of xylem staining, this was not the case in any of the mutants which all had increased fluorescent signal at the inner side of NVE and xylem (Figure 5C and D, Supplementary Figure 9C and D). Thus, the NVE forms an apoplastic barrier which is dependent on expression of *LjMYB36*, *LjSGN1* and *LjSGN3* in a similar manner as in root tissues.

**Figure 5.**
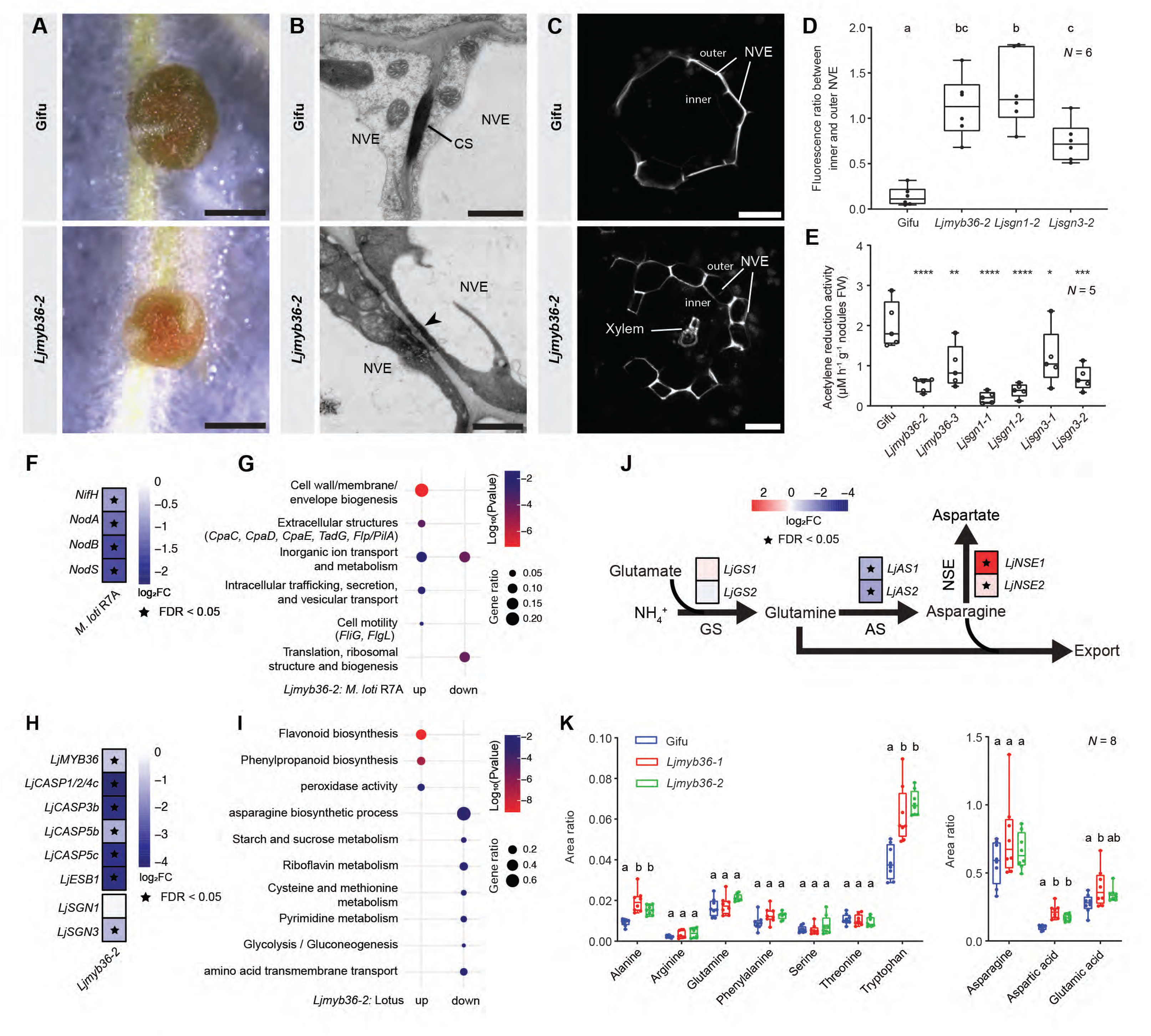
Disturbed nodule function in nodules of Casparian strip defective mutants. (**A**) Representative images of nodules formed at 21 dpi after mass inoculation. (**B**) TEM images of two nodule vascular endodermal cells at 21 dpi. Electron-dense depositions highlighted by line represent Casparian strip (CS) in Gifu nodule vascular endodermis (NVE). Arrowhead indicates the absence of CS in *Ljmyb36-2* NVE. Representative images of at least five nodules from individual plants. (**C**) Nodule vascular endodermis stained with PI dye (white signal). Representative images of at least 10 nodules from individual plants. (**D**) Quantification of fluorescence signal between inner and outer NVE. Different letters depict the statistical difference in a one-way ANOVA analysis with Tukey’s test (*P* < 0.05). (**E**) Quantification of acetylene reduction activity of nodules at 21 dpi. One-way ANOVA analysis with Dunnett’s test was performed between Gifu and mutants (\**P* < 0.05, \*\**P* < 0.01, \*\*\**P* < 0.001, \*\*\*\**P* < 0.0001). (**F**) Heatmap of selected bacterial genes. (**G**) Dotplot of Clusters of Orthologous Group (COG) enriched in bacterial upregulated or downregulated genes. (**H**) Heatmap of host genes involved in CS formation. (**I**) Dotplot of GO terms and KEEG pathways enriched in host DEGs. (**J**) Heatmap of genes involved in converting fixed nitrogen to glutamine, asparagine and aspartate. (**K**) Quantification of amino acids in nodules at 21 dpi. Different letters in depict the statistical difference between Gifu and mutants of a specific amino acid in a one-way ANOVA analysis with Tukey’s test (*P* < 0.05). CS, Casparian strip; NVE, nodule vascular endodermis. Scale bar: 1 mm (**A**), 1 µm (**B**), 20 µm (**C**).

### Functional apoplastic barriers in the nodule vascular endodermis is required for optimized N-fixation

To assess the consequence of dysfunctional CS formation in the NVE on N-fixation, we employed an acetylene reduction assay to measure the nitrogenase activity. When compared to Gifu, mature nodules of all CS mutants showed a significant reduction in nitrogenase activity (Figure 5E). To investigate this in detail, we performed a meta-transcriptome analysis of Gifu and *Ljmyb36-2* nodules. A PCoA showed that the transcriptional outputs of symbiont and host separated according to host genotype (Supplementary Figure 10). In line with the reduced nitrogenase activity, the essential component of nitrogenase *nifH* (*37*) as well as several Nod factor biosynthesis genes were downregulated in rhizobia hosted in *Ljmyb36-2* nodules (Figure 5F). Moreover, Clusters of Orthologous Group (COG) related to “Extracellular structures” and “Cell motility” were enriched in the upregulated genes of bacteria (Fold change > 0, FDR *P*-value < 0.05) of *Ljmyb36-2* (Figure 5G).

Amongst the DEGs from the host, we found genes involved in CS formation (e.g. *LjCASPs*, *LjESB1* and *LjPER64*) to be downregulated in *Ljmyb36-2* nodules (Figure 5H), consistent with the observed absence of CS in NVE. We further found that GO terms and Kyoto Encyclopedia of Genes and Genomes (KEGG) pathways related to several metabolites were repressed in *Ljmyb36-2* nodules, including “asparagine biosynthetic process” and “amino acid transmembrane transport” (Figure 5I). In legume nodules, fixed N in the form of NH_4_^-^ is added to glutamate by the glutamine synthetase enzyme (GS), which can be further converted to asparagine by asparagine synthetases (AS) (*38*) (Figure 5J). Both *LjAS1* and *LjAS2* were downregulated, while *LjNSE1* and *LjNSE2*, which convert asparagine to aspartate, were upregulated in *Ljmyb36-2* nodules (Figure 5J). Thus, despite that we cannot rule out a role of the root CS, the dysfunctional CS in the NVE of *Ljmyb36-2* nodules may result in a shuffling of fixed N into aspartate/aspartic acid. To test this, we performed an LC-MS based analysis of amino acids in the nodules. Indeed, we found that aspartic acid content of homogenized nodules was increased (Figure 5K). Combined, this analysis provides insights into how apoplastic barriers in NVE affect bacterial behavior and nodule homeostasis.

## Discussion

The formation of the CS is a fundamental prerequisite for selective ion uptake in vascular plants. The ability to block out unwanted compounds and provide a filter function is likely a crucial part of sustaining a long-distance transport system such as the vasculature (*1*). Additionally, the formation of symbiotic relationships with microorganisms - arbuscular mycorrhizal fungi - is an important feature which arose even earlier in plant life (*39*). Evidence is emerging that the underlying pathways of barrier establishment are connected to the biotic and nutritional integration of complex soil environments (*18*). Yet, the connection between barrier function and symbiotic relationships has remained elusive. In this work, we have addressed this through a comparison of the network responsible for CS formation in the well-studied Arabidopsis and the symbiosis model species Lotus. Our work illustrate that these species share a conserved machinery in CS formation and that this is required for the establishment and maintenance of a mutualistic inter-species relationship.

This finding agrees with several studies showing that genes controlling CS establishment in Arabidopsis are conserved in rice (*MYB36*, *CASP1, CIF1/2, SGN3*), maize (*ESB-like*) and *Brassica rapa* (*SGN3*) (*12–17*). Intriguingly, this additionally proposes that, in Lotus, the SGN pathway is MYB36-dependent. One outcome of this is that SGN-dependent mechanism such as ectopic lignin deposition and oversuberization does not occur in *Ljmyb36* roots, due to reduced expression of both *LjSGN1* and *LjSGN3*. Similar observations have been made in rice where ectopic suberization only occurs in *Osmyb36a* single mutants (*14*). This could be explained by the fact that both *OsSGN1* and *OsSGN3* are downregulated in *Osmyb36abc*, but not in *Osmyb36a* mutants (*14*). Thus, the MYB36-independent SGN responses may be specific to Arabidopsis. Despite this, previous study revealed that *AtSGN1* and *AtSGN3* are also downregulated in *Atmyb36-2* roots (*18*). This implies that the surveillance system to compensate defective CS is more robust in Arabidopsis, despite the attenuated expression of *AtSGN1/3*. This could be attributable to the anatomy of Arabidopsis roots, which are simpler with only one cortex layer, compared with more complex Lotus, rice and maize roots with multiple layers of cortex. The lack of SGN responses induced by external CIF2 peptides in Lotus roots agrees with a recent study showing that OsCIF2 also cannot activate ectopic lignin deposition in rice (*15*). This suggests that it could be an isolated case in Arabidopsis that CIF peptides alone can activate SGN responses, additional cues are needed in other plants.

The SGN pathway constitutes a linear phosphorylation activated signaling cascade similar to mitogen-activated protein kinases (*40*). Indeed, by employing trans-complementation assay using Lotus *SGN* genes in Arabidopsis *sgn* mutants we were able to show that SGN1 can be activated by a non-native SGN3, whereas the endogenous ligands or endogenous concentration of ligands can only activate native SGN3. Previous biochemical analysis of AtSGN3 revealed that several residues in the receptor ectodomain enable a high-affinity binding of AtCIF2 (*34*). Most of them are conserved in LjSGN3, except in the central binding groove, where only one of three validated residues (Tyr440) is conserved (Supplementary Figure 4C). However, this cannot explain that LjSGN3 is unable to complement the CS fusion-related function in *Atsgn3-3* roots, as mutations of all these three residues in AtSGN3 can still robustly induce CS formation (*34*). Therefore, there must either be other residues different within the LjSGN3 ectodomain, hindering the closure of CS or the configuration of the LjSGN3 in the endodermis is different. In support of the former, in the proposed central binding groove of AtSGN3 (*34*), 9/25 residues are divergent between LjSGN3 and AtSGN3 (Supplementary Figure 4C). Similarly, 6/21 residues are different between LjCIF2 and AtCIF2 mature peptides (Supplementary Figure 2A). As the response of LjSGN3 to exogenous application of native LjCIF2 peptides appears more robust, those species-specific adaptations in both ligand and receptor most likely ensure a high affinity binding. Despite the fact that LjSGN3 cannot restore CS function, it can fully complement the shoot performance of *Atsgn3-3* in the natural soil, and the N-induced growth promotion. This agrees with our previous work that the SGN pathway is required to convey soil condition to affect shoot performance, irrespective of CS status (*18*). Therefore, we propose that SGN3-mediated N signaling is conserved across species, whereas species-specific adaptations of SGN3 with ligands ensure barrier function. This is likely due to yet-to-be-identified additional downstream kinases that are capable of conveying these specific parts of the SGN signaling, or alternatively, converge on ROS signaling via RBOHF activation (*41*).

During nodule formation, endodermal cell divisions are induced at an early primordium stage. To do so, the endodermis has to dedifferentiate to remove CS (*25*). After a few rounds of divisions, CS is reformed at the top cell layer derived from root endodermis (*25*). It was therefore surprising to find that absence of CS has a negative effect on nodule organogenesis. Hence, the absence of CS must have a preexisting effect on nodule formation during the early stages of organogenesis. We provide evidence that CS defective mutants cannot convey N status in the uninfected root to inform shoot to synthesize miR2111. This is a consequence of reduced expression of *CEP1* in the root. However, we cannot completely rule out the possibility that CEP1 peptides can diffuse out of the vasculature in CS defective roots, which may exacerbate the miscommunication between root and shoot. It has been shown that TML represses rhizobial infection and nodule organogenesis (*42*). Therefore, the increased expression of *TML* in CS defective mutant roots could account for the reduced infection and aborted nodule organogenesis. Moreover, we found that auxin and cytokinin homeostasis is disturbed in both *Ljmyb36-2* and *Ljsgn1-2* nodule primordia, whereas exogenous application of auxin or cytokinin can only promote nodulation on *Ljmyb36* roots, but not on *Ljsgn* roots. Thus, the SGN pathway appears to be required for the hormone complementation on CS defective roots.

A recent study showed that *Ljmyb36* mutants have increased sensitive to abiotic stresses, such as salt (*29*). Based on our work, these defects can be due to defective CS establishment. This agrees with several studies showing that disrupted CS formation leads intolerance to abiotic stresses (*16*, *46*). Defect in root apoplastic barrier also affects root exudation (*47*). Therefore, it is interesting to investigate whether the root exudate profile is affected in Lotus CS mutants, and the impact on nodule formation as well as the establishment of other symbiotic relationships, such as arbuscular mycorrhization.

Our work additionally brings forth the idea that CS formation in the endodermis of roots and nodule vascular bundles share a conserved molecular framework. During transition from free-living bacteria to N-fixing bacteroids, COGs related to “Extracellular structures” and “Cell motility” are usually repressed (*48*). However, these COGs were enriched among the upregulated bacterial genes in *Ljmyb36-2,* this proposes a lack of differentiation towards bacteroid stage in the CS mutant nodules. The overaccumulation of aspartic acid in *Ljmyb36* nodules and enriched GO terms related to “amino acid transmembrane transport” in the downregulated DEGs in *Ljmyb36-2* nodules suggest a reduced ability to export amino acids out of mutant nodules. This is supported by the fact that alanine and tryptophan were also over accumulated in *Ljmyb36* nodules. It has been recently shown that barriers in nodule endodermis control oxygen permeation into nodule tissue (*49*). Defective mutants of such barriers have increased oxygen level within nodules and reduced nitrogenase activity. Our work shows that the barriers in nodule vascular endodermis are also important for nodule function, possibly by ensuring fixed N exported out of nodules. This highlights that compartmentalization not only occurs in symbiosome (i.e. intracellular bacteria surrounded by peribacteroid membrane) (*50*), but also in other nodule tissues by forming physical barriers for proper nodule function.

In summary, we demonstrate that Lotus and Arabidopsis share a conserved machinery on CS establishment. More importantly, apoplastic barrier is required for root and shoot communication. By this, a systemic regulation of nodulation is activated, priming the uninfected roots for nodule formation. We also propose that the CS in NVE is important for N-fixation and amino acid homeostasis. Our findings bring insights into the roles of apoplastic barriers in root nodule symbiosis.

## Acknowledgments

The authors thank Ton Timmers and the Central Microscopy facility (CeMic) at MPIPZ for microscopy aid, Aristeidis Stamatakis and the greenhouse team at MPIPZ for help with plant growth. Katharina Markmann and Moritz Sexauer are thanked for sharing of protocols, *tml5* seeds and constructive discussions. Tian Zeng and Caroline Gutjahr are thanked for sharing of protocols. Moreover, we thank Niko Geldner and his lab for insightful discussions. BDR and APM are just thanked. This work was supported by the Sofja Kovalevskaja programme at the Alexander von Humboldt foundation (to TGA) as well as the Max Planck Society (to TGA), German Academic Exchange Service (DAAD, Graduate School Scholarship Program reference number: 91713467 to MM), German Research Foundation (DFG grant MA 7269/1-1 to MM), the Cluster of Excellence for Plant Sciences (CEPLAS) under Germany’s Excellence Strategy EXC-2048/1 (project ID 390686111 to NDR and SMe).

## Author contributions

Conceptualization: DS and TGA. Methodology: DS, REV, UN, NDR, SMe, RTN, MM, TGA. Investigation: DS, REV, UN, NDR, SMa, SMe, RTN. Visualization: DS and TGA. Funding acquisition: TGA. Main writing: DS and TGA. All authors read and commented on the final version of the manuscript.

## Competing interests

All authors declare that they have no competing interests.

## Data and materials availability

RNA-seq raw reads generated in this study have been deposited at National Center for Biotechnology Information under BioProject ID PRJNA1022778 and PRJNA1022988.

## Materials and Methods

### Plant materials and growth conditions

*Lotus japonicus* (Lotus) seeds were sandpaper scarified and surface sterilized with 1% sodium hypochlorite for 10 min, then washed six times and incubated on a shaker in sterile ddH_2_O at room temperature for 1 h. Seeds were transferred to petri dishes (IZl 90 mm) with wet filter paper and stratified at 4°C in darkness for 2-3 days. Seeds were germinated at 21°C in darkness for two days, then transferred to square plates (12×12cm) containing 40 mL of ¼ B&D medium (*51*), supplemented with 0.5 mM KNO_3_ at 21°C under a 16-h-light/8-h-dark photoperiod. For Lotus growth, ¼ B&D medium was always supplemented with 0.5 mM KNO_3_. For CIF2 peptide treatment on agar medium, 6-day-old Lotus seedlings were grown as above-mentioned, transferred to square plates containing 40 mL of ¼ B&D agar plates supplemented with 500 nM peptide or H_2_O (mock), and grown for 2 days. For CIF2 peptide treatment on hydroponics system, 2-day-old germinated seedlings were transferred to open and sterile WECK jar filled with glass beads (IZl 5 mm) and ¼ B&D liquid medium. 6-day-old seedlings were treated with 500 nM peptide or H_2_O (mock) for 2 days after refreshing the liquid medium. For LjCEP1 peptide treatment on agar medium, 2-day-old germinated seedlings were transferred to square plates containing 25 mL of ¼ B&D agar plates supplemented with 1 µM LjCEP1 peptides or H_2_O (mock). Note that medium volume per square plate was reduced compared with plates used for other nodulation assays to save LjCEP1. For CEP1 peptide treatment on plants grown in a leca:vermiculite mixture (4:1), 2-day-old germinated seedlings were transferred to mixture soaked with ¼ B&D liquid medium, and grown for 5 days. 0.1% Triton X-100 solution containing 1 µM LjCEP1 peptides or H_2_O (mock) was directly applied onto the leaves of 7-day-old plants, ∼ 20 µL per plant, three times per week, for two weeks. To create *Ljmyb36-2xsgn3-2, Ljmyb36-2xtml5* and *Ljsgn3-2xtml5* double mutants, emasculated *Ljmyb36-2* was pollinated by *Ljsgn3-2* or *tml5*, emasculated *Ljsgn3-2* was pollinated by *tml5.* Homozygous double mutants were identified in the F2 population. Lotus LORE1 mutants were genotyped following CTAB-based DNA isolation, primers used to genotype Lotus *LORE1* mutants and *LORE1* mutant lines used in this study are listed in Supplementary Table 3.

*Arabidopsis thaliana* (Arabidopsis) seeds were sterilized by incubation in a solution of 70% ethanol and 0.05% Triton X-100 for 5 min. Seeds were briefly washed with 100% ethanol, and dried on filter paper. Seeds were then evenly poured on square plates containing ½ Murashige and Skoog (MS) medium, stratified at 4°C in darkness for 2-3 days, and grown at 21°C under a 16-h-light/8-h-dark photoperiod. For CIF2 peptide treatment on agar medium, 5-day-old seedlings grown on ½ MS were transferred to square plates containing 40 mL of ½ MS agar, supplemented with 100 nM peptide or H_2_O (mock), and grown for 2 days. For Arabidopsis plants grown on CAS, 7-day-old seedlings grown on ½ MS were transferred to square pots (9×9 cm) containing CAS. Pots were placed on top of a capillary mattress in a tray, and regularly watered with tap water or 7% calcinit^TM^ solution (0.651 g/L, calcinit^TM^ composition: 14.4% nitrate, 1% ammonium, 26% calcium oxide) (Yara, Germany) from the bottom.

### Cloning

Constructs for transforming Lotus were all generated using the Golden Gate system (New England BioLabs) (*52*). Promoter regions of *LjSCR* (3137 bp), *LjMYB36* (3800 bp), *LjSGN1* (2749 bp), *LjSGN3* (3123 bp) and *LjSHR* (3040 bp) were cloned into pICH41295. Gene sequences of *LjMYB36* (1444 bp, introns inclusive), coding sequence of *LjSGN1* (1281 bp), *LjSGN3* (3819 bp) and *LjCEP1* (378 bp) were cloned into pICH41308. 3’UTR of *LjMYB36* (509 bp), *LjSGN1* (595 bp), *LjSGN3* (530 bp) and *LjSHR* (1066 bp) were cloned into pICH41276. pICH47751 and pICH47761 were used as the level 1 acceptors to assemble level 0 vectors. *pLjSCR:LjSGN1cds:act2Ter* (pICH44300), *pLjSCR:β-glucuronidase* (*GUS*)(pICH75111)*:act2Ter*, *pLjMYB36:LjMYB36gene:LjMYB36 3UTR*, *pLjSGN1:LjSGN1cds: LjSGN1 3UTR*, *pLjSGN3:LjSGN3cds:LjSGN3 3UTR*, *pLjMYB36:GUS:LjMYB36 3UTR*, *pLjSGN1:GUS:LjSGN1 3UTR*, *pLjSGN3:GUS:LjSGN3 3UTR*, *pLjSHR:LjCEP1cds:LjSHR 3UTR, pLjSHR:GUS:LjSHR 3UTR* were assembled using pICH47751. *pLjSCR:LjSGN3cds:act2Ter* was assembled in pICH47761 for overexpressing *LjSGN3* together with *LjSGN1*. *pAtUBQ10:NLS 3xmScarlet:nosTer* (pICH41421) were assembled using pICH47732 for detecting transgenic roots, pICSL70004 expressing *pNos:NPTIIcds:OcsTer* was cloned into pICH47742 for selecting transgenic roots. pAGM4673 was used as the level 2 acceptor to assemble level 1 vectors, and transformed to *Agrobacterium rhizogenes* MSU440 by electroporation.

Constructs for transformation Arabidopsis were all generated using the Gateway system (Invitrogen). Coding sequences of *LjSGN1* (1281 bp) and *LjSGN3* (3819 bp) were cloned into pDONR221. For the chimeric version, N-terminal region of AtSGN3 CDS (1-1260 bp) and C-terminal region of LjSGN3 CDS (2659-3816 bp) were merged by PCR. For subcellular localization of SGN3, coding sequence of mTurquoise2 was added at the C-terminal region of AtSGN3, LjSGN3 or chimeric AtLjSGN3 without stop codon by PCR, and cloned into pDONR221. pAtSGN1, pAtSGN3, AtSGN1 3UTR and AtSGN3 3UTR (*11*, *43*), together with generated pDONR221 vectors were assembled using the binary vector pED97 (*53*). Binary vectors were transformed into *sgn1*-2 or *sgn3*-3 expressing *AtCASP1pro:AtCASP1-GFP* (*11*, *43*) using the floral dip method (*54*). Transformed seeds expressing FastRed were selected using a Zeiss Axio Zoom.V16 stereomicroscope. Two independent T3 homogenous lines were used for analysis. Primers used for cloning are listed in Supplementary Table 3.

### Staining procedures and microscopy

All staining procedures for whole-mount roots were done using ClearSee staining (*55*, *56*). Briefly, roots were fixed in 1 x PBS containing 4% paraformaldehyde (w/v) at 4°C overnight, following vacuum treatment for 30 min. Samples were washed three times with 1 x PBS, and cleared using ClearSee solution (10% xylitol, 15% sodium deoxycholate and 25% urea in water) till sample became clear under gentle shaking. Samples were transferred to new ClearSee solution containing 0.2% Basic Fuchsin for lignin staining. The dye solution was removed after overnight staining and washed with fresh ClearSee solution overnight till no red coloration. Fluorol Yellow 088 (Santa Cruz Biotechnology)(FY) staining of Arabidopsis and Lotus roots for suberization pattern was performed as previously described (*43*, *56*). Suberization pattern in Arabidopsis roots were determined as previously described (*57*). Suberization pattern in Lotus roots was determined as following: start of patchy zone, first suberized endodermis cell (usually associated with phloem poles); start of continuously suberized zone: start of continuous endodermis cells associated with phloem poles are suberized. See Supplementary Figure 1F for an example. Note that in contrast to Arabidopsis, the endodermis cells associated with xylem poles are not continuously suberized in Lotus roots. For propidium iodide (PI) assays on Lotus roots, whole seedlings were incubated in 0.01% (w/v) PI solution (Sigma) for 10 min, and rinsed in water for 2-3 min. For PI assays on mature nodules, whole nodules with attached roots (∼2 mm) were incubated in 0.01% (w/v) PI solution at 4°C overnight, washed with water at room temperature for at least three times, each wash for 1 h. Cross-sections on PI-stained nodules at the middle position were made by hands. PI-stained samples were transferred to chambered slides with cover slips for imaging. Confocal images of Basic Fuchsin staining were taken using a Leica SP8 FALCON-DIVE confocal microscope for Lotus, and using a Zeiss LSM 980 system for Arabidopsis. Fluorophore settings were ex 1055 nm, em 600-650 nm for Lotus; ex 561 nm, em 600-650 nm for Arabidopsis. Confocal images of PI penetration and FY staining for Lotus roots were taken using a Zeiss LSM 980 system. Confocal images of PI-stained nodules were taken using a Zeiss LSM 880 system. Fluorophore settings were ex 514 nm, em 600-650 nm for PI; ex 488nm, em 500-550 nm for FY. FY staining for Arabidopsis roots were imaged using a Zeiss Axio Zoom.V16 stereomicroscope equipped with a Zeiss Axiocam 705 mono or color camera. Software ImageJ was used to analyze fluorescence intensity of PI signal on NVE (convert to 8-bit, plot profile over a fixed 25-micron line marked across a representative NVE, after straighten over 20 pixels), numbers on inner and outer NVE position were used to calculate fluorescence signal between inner and outer NVE.

For semi-thin sections on nodule primordia and nodules, samples were fixed in 4% (w/v) paraformaldehyde, 2.5% (v/v) glutaraldehyde in 0.05 M sodium phosphate buffer (pH 7.2) at 4°C overnight. The fixed samples were dehydrated in an ethanol series and embedded in Technovit 7100 (Heraeus Kulzer) according to the manufacturer’s protocol. Sections (5 to 12 mm) were cut using a Leitz 1512 microtome and stained for 1.5 min in 0.05% (w/v) toluidine blue O or 10 min in 0.1% (w/v) ruthenium red to examine transgenic GUS material. The sections were imaged using a Zeiss Axio Imager.A2 microscope equipped with a Zeiss Axiocam HRc camera.

### Nodulation assays

For nodulation assays on Lotus, *Mesorhizobium loti* R7A was used unless otherwise stated. Rhizobia liquid cultures were grown at 28°C for 2 days and pelleted for 5 min at 3000 g. The bacterial pellet was resuspended in sterile ddH_2_O and adjusted to OD_600_ = 0.01 with sterile ddH_2_O. For spot inoculation assay, ∼0.5 µl bacterial suspension was applied on the root susceptible zone under a stereomicroscope. For mass inoculation assay on plates, 500 µl bacterial suspension was applied per plate. For nodulation assays on plates, inoculations were performed at 7 days after germination (2 days of germination in darkness inclusive). For nodulation assays on transformed Lotus roots grown in a leca:vermiculite mixture (4:1), 1 mL bacterial suspension was applied per plant at one week after transplanting. For nodulation assays on LjCEP1-treated Lotus grown in mixture, 1 mL bacterial suspension was applied per plant at 9 days after transplanting. Nodule primordium and nodule numbers were counted under a stereomicroscope, and/or harvested at the time points mentioned in the text. For visualizing infection threads, roots inoculated with *M. loti* LacZ strain were fixed with 1.25% (v/v) glutaraldehyde in 200 mM sodium phosphate (pH 7.0) at room temperature under vacuum for 1 h, then rinsed three times with 200 mM sodium phosphate (pH 7.0). Samples were incubated under vacuum for 1 h in 200 mM sodium phosphate (pH 7.0) containing 0.08% (w/v) X-Gal (5-bromo-4-chloro-3-indolyl-β-D-galactopyranoside), 5 mM K_3_(Fe(CN)_6_) and 5 mM K_4_(Fe(CN)_6_). This was followed by an overnight incubation under atmospheric pressure in darkness at 28°C. Samples were then rinsed three times with 200 mM sodium phosphate buffer (pH 7.0), and finally observed for infection threads under a light microscope.

### Transcriptomics

For transcriptomic analysis on roots, whole roots of 8-day-old seedlings grown on ¼ BD agar plates were harvested, ∼10 roots per biological replicate. For the time-course transcriptomic analysis on nodule primordia, 7-day-old Gifu, *Ljmyb36-2*, *Ljsgn1*-2 seedlings were spot inoculated at the susceptible zone, positions of inoculums were marked on filter paper. Root segments (∼3 mm) at the marked positions were harvested at 0, 1, 3, and 5 days post-inoculation (dpi), ∼15 root segments per biological replicate. For metatranscriptomic analysis on nodules, 21-day-old Gifu and *Ljmyb36-2* nodules were harvested, one nodule per plant, ∼10 nodules per biological replicate. Note that only dark-colored nodules of Gifu and *Ljmyb36-2* were harvested. Samples were all immediately frozen in liquid nitrogen. RNA of roots and nodule primordia was extracted using a TRIzol (Invitrogen)-adapted ReliaPrep RNA extraction kit (Promega), as previously described (*53*). RNA of nodules was extracted using a RNeasy Plant Mini Kit (Qiagen) according to the manufacturer’s manual. RNA quality was determined using a Bioanalyzer 2100 system (Agilent Technologies). Library preparation and paired-end 150 bp sequencing were conducted by Novogene (Cambridge, UK). Approximately 40 million raw reads were generated per sample. RNA-seq raw reads of Col-0, *Atmyb36-2*, *Atsgn3-3 and Atmyb36-2xsgn3-3* roots were acquired from (*32*). All raw reads were preprocessed using fastp (v0.22.0) (*58*). Filtered high-quality reads were mapped to A. thaliana TAIR10 reference genome with Araport 11 annotation (Phytozome genome ID: 447) or *L. japonicus* Gifu v1.2 genome assembly (*59*) with Gifu v1.3 annotation (https://lotus.au.dk), or *M. loti* R7A genome with annotation (JGI Project ID: 1078770) with using HISAT2 (v2.2.1) (*60*), and counted using featureCounts from the Subread package (v2.0.1) (*61*). All statistical analyses were performed using R (v4.1.2) (https://www.R-project.org/). Raw read counts were normalized using TMM normalization (calcNormFactors), and transformed to counts-per-million (cpm) using edgeR package (*62*). Lowly expressed (less than 5 cpm all samples combined, and at least 1 cpm in 2 samples) genes were removed from the analysis. Differentially expressed genes (DEGs) were identified by fitting a negative binomial generalized linear model to the genes using the glmFit function in edgeR package (*62*), with absolute fold change higher than 2 and a false discovery rate (FDR) corrected *P*-value lower than 0.05. For Gene Ontology (GO) term enrichment and KEGG pathway analysis, Gifu v1.3 protein sequences were annotated using eggNOG-mapper v2 (eggNOG 5 database, default annotation options) (*63*). Note that GO term annotations not related to plants were removed. GO term enrichment analysis on DEGs was performed using Metascape (*64*) for Arabidopsis, topGO package for Lotus (https://bioconductor.org/packages/topGO). KEGG pathway and Clusters of orthologous groups (COGs) enrichment analysis were performed using clusterProfiler package (*65*). Heatmaps were generated using pheatmap package (https://cran.r-project.org/web/packages/pheatmap/index.html) or ggplot2 package (*66*). PCoA was performed using ggplot2 package (*66*). No custom scripts were generated in this study. DEGs of all transcriptome analysis are listed in Supplementary Table 4.

### Hairy root transformation

Transient root transformation was conducted by the hairy root method (*67*). 7-day-old Lotus seedlings grown on Gamborg’s B5 agar plates were cut and hypocotyls were submerged in an *A. rhizogenes* MSU440 suspension transformed with constructs. Treated hypocotyls were placed on Gamborg’s B5 agar and incubated at 21°C in darkness for 2 days. Hypocotyls were then incubated at 21°C under a 16-h-light/8-h-dark photoperiod for 3 days. Hypocotyls were transferred to Gamborg’s B5 agar plates supplemented with cefotaxime (300 μg/mL) to remove *A. rhizogenes* and with neomycin (25 μg/mL) to inhibit the growth of untransformed roots. Composite plants for axenic root development analysis were transferred to ¼ B&D agar plates, and grown for 7 days. Composite plants for nodulation assay were screened for the presence of an NLS-3xmScarlet transformation marker under a Zeiss Axio Zoom.V16 equipped with a red filter. Plants bearing transformed roots were transferred to a sterile leca:vermiculite mixture (4:1) soaked with ¼ BD liquid medium. Roots or nodules expressing promoter:GUS constructs were incubated in staining buffer containing 0.05% (w/v) of X-Gluc (5-Bromo-4-chloro-1H-indol-3-yl β-D-glucopyranosiduronic acid), 100 mM phosphate buffer (pH 7.0), 0.5 μM EDTA (pH 7.0), 0.5 mM K_3_(Fe(CN)_6_) and 0.5 mM K_4_(Fe(CN)_6_) under vacuum treatment at room temperature for 15 min. Samples were then incubated in the staining buffer at 37°C till GUS signals were visible. Samples were then washed three times with 100 mM phosphate buffer (pH 7.0) before imaging. Overviews of roots and nodules were imaged using a Zeiss Axio Zoom.V16 stereomicroscope equipped with a Zeiss Axiocam 705 color camera.

### Quantitative RT-PCR

LjCEP1 or mock treated Lotus plants were grown as above-mentioned. Tissues of 16-day-old seedlings were harvested, ∼8 shoots, roots or susceptible zones (∼3 mm) per biological replicate, three to four biological replicates per sample. Samples were immediately frozen in liquid nitrogen. RNA extraction and cDNA synthesis were performed as previously described (*23*). cDNA synthesis was performed using the RevertAid First Strand cDNA Synthesis Kit (Thermo Fisher) in a final volume of 20 μL. Each reaction contained 1 μg of total RNA (for shoots and roots) or 200 ng of total RNA (for susceptible zones), 1 μL of 1 mM Oligo(dT)_18_, and primers targeting LjU6 and miR2111, 0.5 μL of 1 mM each. qRT-PCR was performed in a BioRad CFX Connect Real-Time system in a final volume of 10 μL. Each reaction contained 5 μL of 2 x iQ SYBR Green supermix (Bio-Rad), 2 μL of diluted cDNA (10 times dilution for shoots and roots, 2 times dilution for susceptible zones with water), 1 µl of 2.5 μM forward primer, 1 µl of 2.5 μM reverse primer and 1 µl of water. LjU6 was used as the reference for miR2111, *LjPP2A* and *LjATPase* were used as the references for other targets. The thermal cycler conditions were: 95 °C for 2min, 40 cycles of 95°C for 30 s, 60°C for 30 s, and 72°C for 30 s. For the melting curve, conditions were set as: denaturation, 95°C for 10 s; hybridization, 60°C for 5 s; denaturation until 95°C with 0.5°C incrementation. Relative expression values were determined using the 2^-ΔΔCT^ method. qRT-PCR primers used in this study are listed in Supplementary Table 3.

### Transmission electron microscopy

For visualizing Casparian strips in nodule vascular endodermis by transmission electron microscopy, 500 µm slices of 21-day-old nodules of Gifu, *Ljmyb36-2*, *Ljsgn1-2* and *Ljsgn3-2* derived from agar plates were cut from the middle of the nodule with a razor blade, perpendicularly to the nodule vasculature. Nodule slices were placed into primary fixative (2% formaldehyde, 2.5% glutaraldehyde in 0.05 M sodium cacodylate buffer pH 6.9) and left with gentle agitation for 2-3 h at room temperature, then at 4°C overnight. After rinsing in 0.05 M sodium cacodylate buffer (4 x 15 min), samples were post-fixed in 1% osmium tetroxide and 1.5% potassium ferrocyanide in 0.05 M sodium cacodylate buffer pH 6.9 for one hour at room temperature and were subsequently rinsed with water. Samples were further dehydrated with a series of ethanol, gradually transferred to acetone and embedded into Araldite 502/Embed 812 resin (EMS; catalog no. 13940) over five days using the ultrarapid infiltration by centrifugation method revisited by McDonald (2014) for the pure resin steps on the last day. Ultrathin (70–90 nm) sections were collected on nickel slot grids as described by Moran and Rowley (1987), stained with 0.1% potassium permanganate in 0.1 N H_2_SO_4_ (Sawaguchi et al., 2001), followed by staining with uranyl acetate replacement stain (UAR-EMS; catalog no. 22405), and lead citrate (Science Services; catalog no. DM22410) for one minute each. Sections were examined with a Hitachi HT7800 TEM (Hitachi High-Technologies Europe) operating at 100 kV and fitted with an EMSIS XAROSA digital camera (EMSIS GmbH Münster).

### Acetylene Reduction Assay

Nitrogenase activity was measured by the reduction of acetylene (C_2_H_2_) into ethylene (C_2_H_4_) and detected using Gas Chromatography - Flame Ionization Detection. Nodulated Gifu*, Ljmyb36-2*, *Ljmyb36-3*, *Ljsgn1-1*, *Ljsgn1-2*, *Ljsgn3-1* and *Ljsgn3-2* plants grown on agar plates, as previously described, were assayed at 21 dpi. Five biological replicates were examined for each line, with a single replicate consisting of two plants. Plants were placed in a 25 mL glass tube with 500 μL of FAB medium (500 μM MgSO_4_·7H2O; 250μM KH_2_PO_4_; 250 μM KCl; 250 μM CaCl_2_·2H_2_O; 100 μM KNO_3_;25 μm Fe-EDDHA; 50 μM H_3_BO_3_;25 μM MnSO_4_·H_2_O;10 μM ZnSO_4_·7H_2_O; 0.5 μM Na_2_MoO_4_·2H2O; 0.2 μM CuSO_4_·5H_2_O; 0.2 μM CoCl_2_·6H_2_O; pH 5.7) (*68*) and sealed with a rubber stopper. Then, 1 mL of air was withdrawn and substituted with 1 mL of C_2_H_2_. Subsequently, 1 mL of the resulting mixture was introduced into a GC 2010 Pro (Shimadzu). Measurements were taken at five distinct time points: 0, 20, 40, 60, and 80 min, with the samples maintained at 28°C in a water bath.

Regression analysis was employed to convert the area under the curve into nanomoles of C_2_H_4_, referencing a C_2_H_4_ standard curve. The nanomoles of C_2_H_4_ per hour were calculated by multiplying the slope of a linear regression model (*C_2_H_4_ nanomoles* = *mt* + *b*, where *m*: slope, *t*: time, and *b*: intercept) by 1 h (60 min).

### LC-MS analysis

Amino acid levels were determined using the Dionex UltiMate 3000 UPLC system (Thermo Scientific, Germering, Germany) coupled to a maXis 4G (Bruker Daltonics, Bremen, Germany) quadrupole-time-of-flight (Q-TOF) mass spectrometer outfitted with an electrospray ion source. Nodules of Gifu, *Ljmyb36-2* and *Ljmyb36-3* plants grown on vermiculite mixture were sampled at 21 dpi and snap frozen. Plant material was ground under liquid nitrogen in sample tubes using a plastic pestle, and aliquots of 20 mg of dry weight were extracted in 1 mL of 80% (v/v) ice-cold aqueous methanol by promptly vortexing and subsequent thermal incubation (800 rpm, 80°C, 15 min) in the ThermoMixer® C (Eppendorf, Hamburg, Germany). After centrifugation (16000 x g, 4°C, 5 min), the supernatant was collected and the pelleted material was re-extracted using 50% (v/v) ice-cold aqueous methanol as mentioned before. The resulting polar supernatant was combined with the former apolar for LC-MS analysis. Separation was performed on a XSelect® CSH (3.5 µm, 2.1×150 mm) column (Waters, Drinagh, Ireland) at room temperature within a 30 min gradient at a flow rate of 0.35 ml/min using a binary gradient system with water/0.1% formic acid (FA) as mobile phase A and methanol/0.1% FA as mobile phase B. The following gradient was used: 0-1 min 2% B (v/v), 1-18 min linear gradient to 99% B, 18-22 min 99% B, 22-23 min linear gradient to 2% B and 23-30 min 2% B. The amino acids were analyzed in negative-ion mode with following source settings: capillary voltage: 3.5 kV, nebulizer pressure: 1 bar, dry gas flow: 8 l/min, dry temperature: 200°C. Data acquisition was carried out under control of the Compass HyStar software (version 6.0) (Bruker, Bremen, Germany). Amino acids were quantified from full-scan MS data (mass range 50-1000 m/z) using the DataAnalysis and QuantAnalysis (both version 6.0) software (Bruker, Bremen, Germany).

### Peptides

AtCIF2 peptides [DY(sulphated)GHSSPKPKLVRPPFKLIPN] and LjCEP1 peptides [AFEP(hydroxylated)TTPGNSP(hydroxylated)GVGH] were synthesized by Pepmic Co., Ltd (Suzhou, China). LjCIF2_sTyr2_ peptides [DY(sulfated)GRYDPTPKLSKPPFKLIPN] and LjCIF2_sTyr2Tyr5_ peptides [DY(sulfated)GRY(sulfated)DPTPKLSKPPFKLIPN] were synthesized by GL Biochem Ltd (Shanghai, China).

### Anthocyanin content measurement

Measurement of anthocyanin content in Arabidopsis rosettes grown under CAS conditions was performed as previously described (*69*). Absorbances at 530 and 637 nm were measured using a Tecan Infinite 200 PRO plate reader.

### Phylogenetic Analysis

For Lotus and *Medicago truncatula* protein sequences, a local BLASTP was performed against Gifu v1.3 protein and *M. truncatula* A17 r5.0 protein sequences (cut-off Expect (E) threshold < −5). Protein sequences of other species were retrieved from Phytozome v13 by BLASTP using Arabidopsis protein sequences (cut-off Expect (E) threshold < −5). Full-length protein sequences were aligned using MAFFT v7.450 (*70*) with default parameter settings (auto algorithm; scoring matrix, BLOSUM62; gap opening penalty, 1.53; offset value: 0.123). Curated alignment was used for tree building using FastTree v2.1.11 (*71*) with the default parameter settings. Branch support analysis was performed using the Shimodaira-Hasegawa test on the three alternate topologies (NNIs) around that split, based on 1000 resamples. Only branches containing genes of interested and their closest homologs were sampled again, aligned and a final phylogeny tree was built as above-mentioned. For the inference of other orthologues (e.g. CASPs, ESB1, PER64, YUCCAs, PINs) in Lotus, each phylogenetic tree was built as above-mentioned. For the phylogenetic tree of CIF1 and CIF2, Lotus CIF1 and CIF2 protein sequences were not annotated in Gifu v1.3, therefore retrieved from MG20 v3.0 by a local BLASP. A phylogenic tree on full-length protein sequences was built as above-mentioned, only the regions encoding mature CIF1 and CIF2 peptides were shown in Supplementary Figure 2A.

### Acronyms and accession numbers

All acronyms with accession numbers used in this study are listed in Supplementary Table 5.

---

**Supplementary Figure 1.**
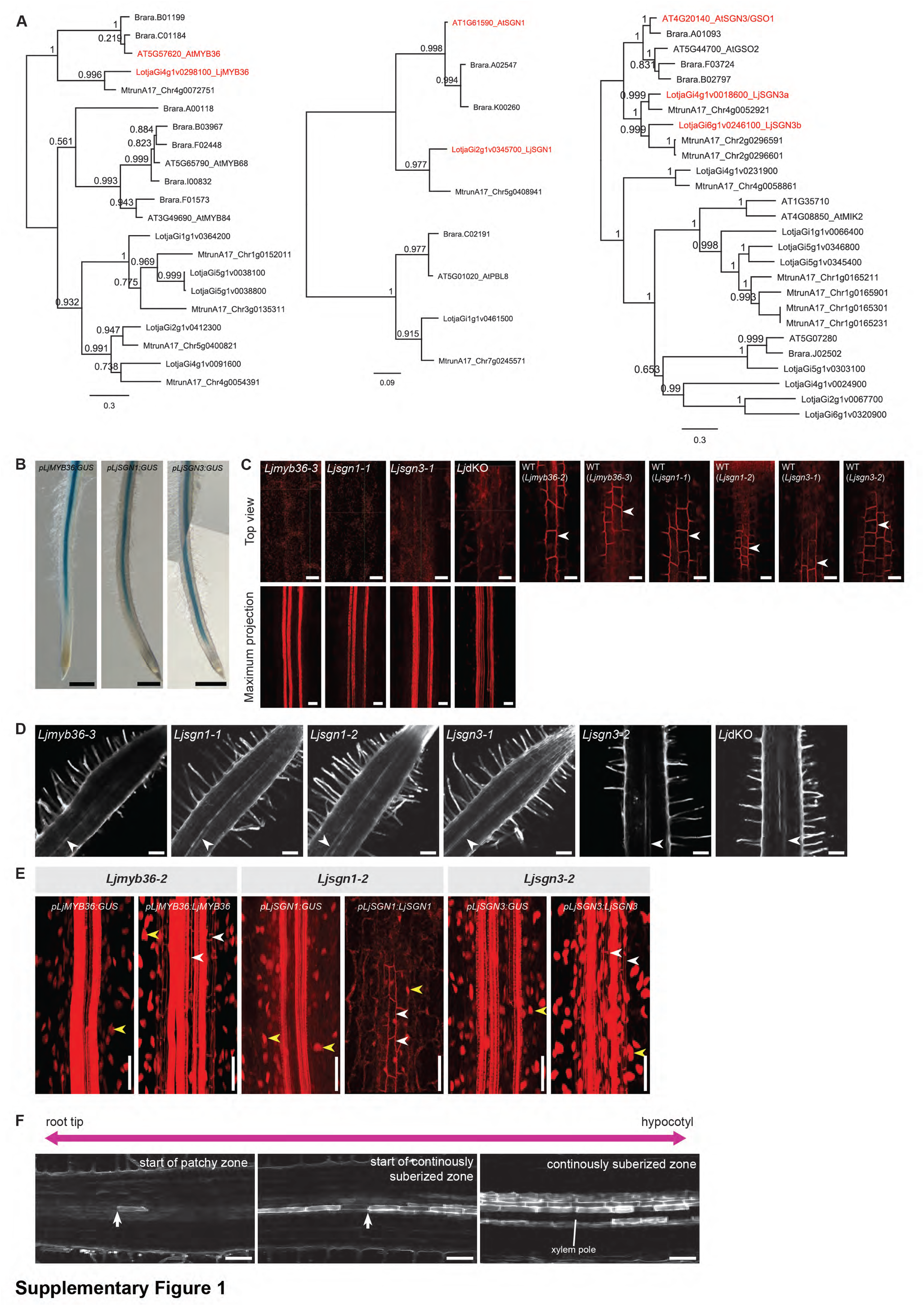
(**A**) Maximum likelihood phylogenetic tree of MYB36 (left), SGN1 (middle) and SGN3 (right) in Arabidopsis, Lotus and other species. The branch containing the closest homolog(s) in Arabidopsis, *Brassica rapa*, *Medicago truncatula* and Lotus was used as the outgroup. Numbers at the nodes indicate FastTree support values. Branch lengths were scaled (see the bottom) and represent the number of substitutions per site. (**B**) *pLjMYB36:GUS, pLjSGN1:GUS* and *pLjSGN3:GUS* activity in Gifu hairy roots. Representative images of at least 10 composite plants. (**C**) Top view (upper) and Maximum projection (lower) of confocal image stacks of Basic Fuchsin stained and fixed 9-day-old roots in the region where CS is formed. Arrowhead indicates CS. Representative images from three independent experiments (n=18) for mutants, one independent experiment for segregated wild-type (WT) lines (n>5). (**D**) Representative images of propidium Iodide (PI) stained 9-day-old roots. Arrowheads indicate no blockage of PI penetration into vasculature. Representative images from two independent experiments. (**E**) Basic Fuchsin staining of complemented mutant roots. White arrowhead indicates Casparian strip (CS), yellow arrowheads indicate transformed root marker of NLS-3xmScarlet. Representative images of at least 6 composite plants. (**F**) Representation of suberization pattern in Fluorol Yellow stained Lotus roots. Arrows indicate start of patchy zone (left) and start of continuously suberized zone (middle). Note that in the continuously suberized zone, only endodermis associated with phloem poles are continuously suberized, endodermis associated with xylem poles (white line) are not continuously suberized. Scale bar: 500 µm (**B**), 20 µm (**C**), 100 µm (**D**), 50 µm (**E**), 1 mm (**F**).

**Supplementary Figure 2.**
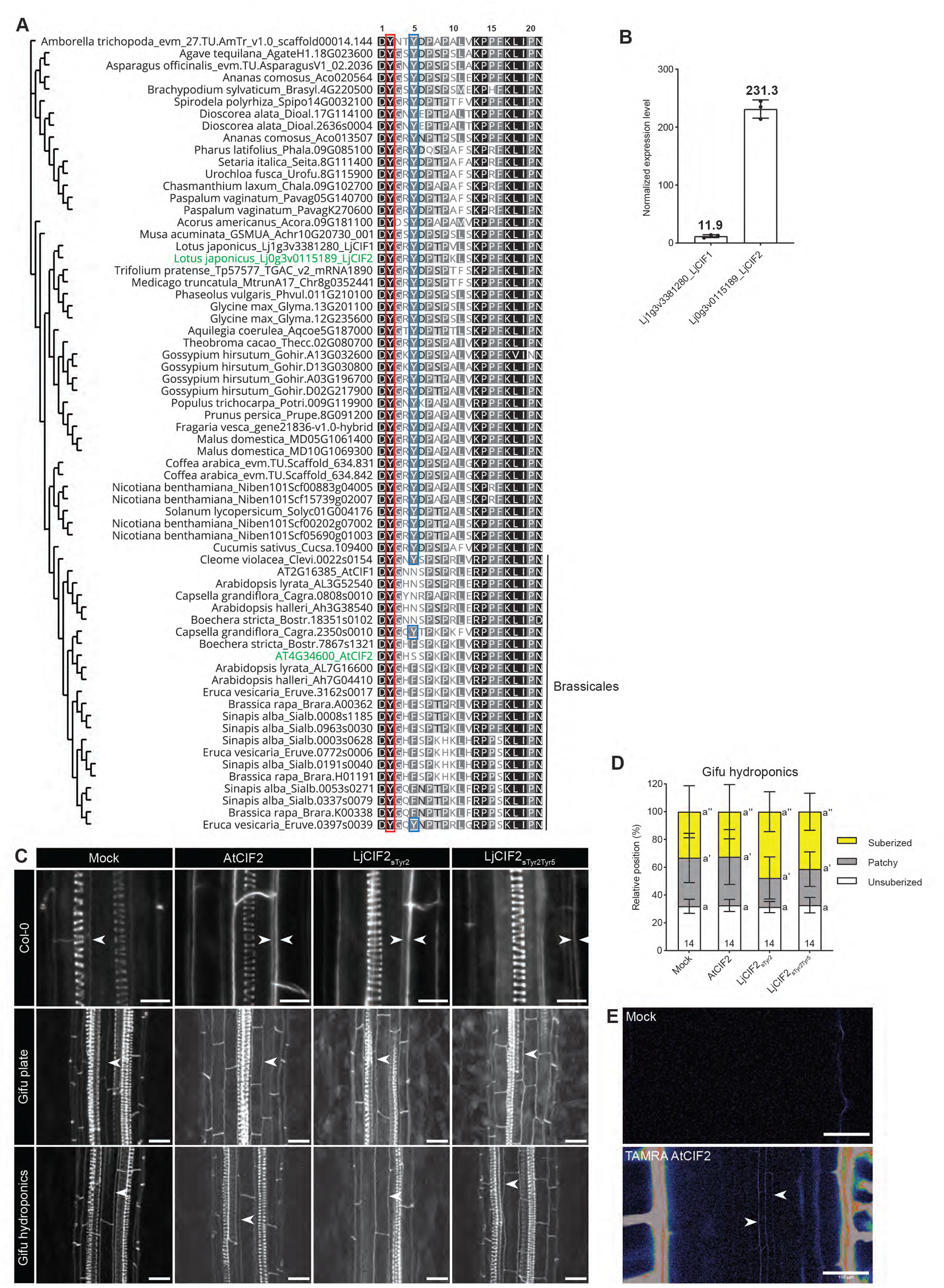
(**A**) Maximum likelihood phylogenetic tree of CIF1 and CIF2 in plant species of angiosperms based on full-length protein sequences. As *Amborella trichopoda* is the most basal angiosperm in the analysis, *Amborella trichopoda_evm_27.TU.AmTr_v1.0_scaffold00014.144* was used as the outgroup. Only mature peptide sequences are shown here. Red box indicates conserved Tyr at position 2 across angiosperm. Blue box indicates conserved Tyr at position 5 across angiosperm, but rarely in Brassicales species. AtCIF2 and LjCIF2 are highlighted in green. (**B**) Normalized expression values of *LjCIF1* and *LjCIF2* in Lotus roots, data from Suzaki T et al., of Lotus Base. Mean values shown on bar. (**C**) Max projection of confocal image stack of Basic Fuchsin stained Arabidopsis and Lotus roots treated with CIF2 peptides. Double arrowheads indicate overlignification. (**D**) Suberization pattern of 8-day-old Lotus Gifu roots after 48 h treatment with 500 nM CIF2 peptide under hydroponic condition. Different letters depict the statistical difference in a one-way ANOVA analysis with Tukey’s test (*P* < 0.05). Representative results from two independent experiments. (**E**) Confocal image of 8-day-old Lotus Gifu roots after 2 h treatment with 500 nM TAMRA-tagged AtCIF2 peptide under hydroponic condition. Arrowheads indicate penetration of TAMRA-tagged AtCIF2 to endodermis at root differentiation zone. Representative image of three roots. Scale bar: 10 µm for Col-0, 20 µm for Gifu (**C**); 100 µm (**E**).

**Supplementary Figure 3.**
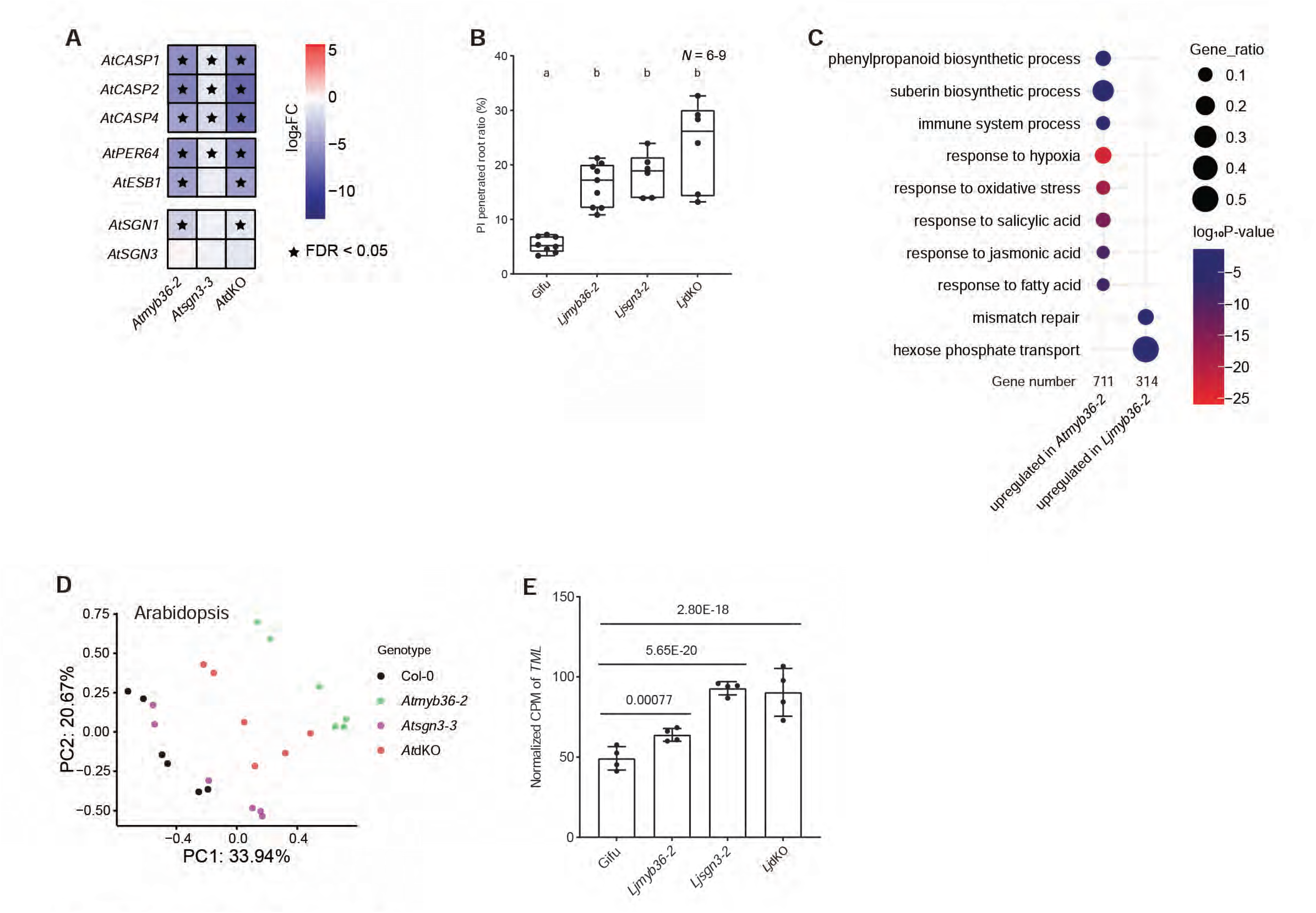
(**A**) Heatmap of genes involved in Casparian strip (CS) formation in Arabidopsis mutants (*32*). (**B**) Quantification of the proportion of 9-day-old roots that can be penetrated by propidium iodide (PI) into the vascular bundle. Representative results from two independent experiments. Different letters depict the statistical difference in a one-way ANOVA analysis with Tukey’s test (*P* < 0.05). (**C**) Dotplot of Gene Ontology terms enriched in upregulated differentially expressed genes in *Atmyb36-2* (*32*) and *Ljmyb36-2* roots. (**D**) Principal coordinate analysis (PCoA) plot of root transcriptome of Arabidopsis CS mutant roots. (**E**) Normalized CPM of *TML* in Lotus CS mutant roots. FDR corrected *P* values are indicated.

**Supplementary Figure 4.**
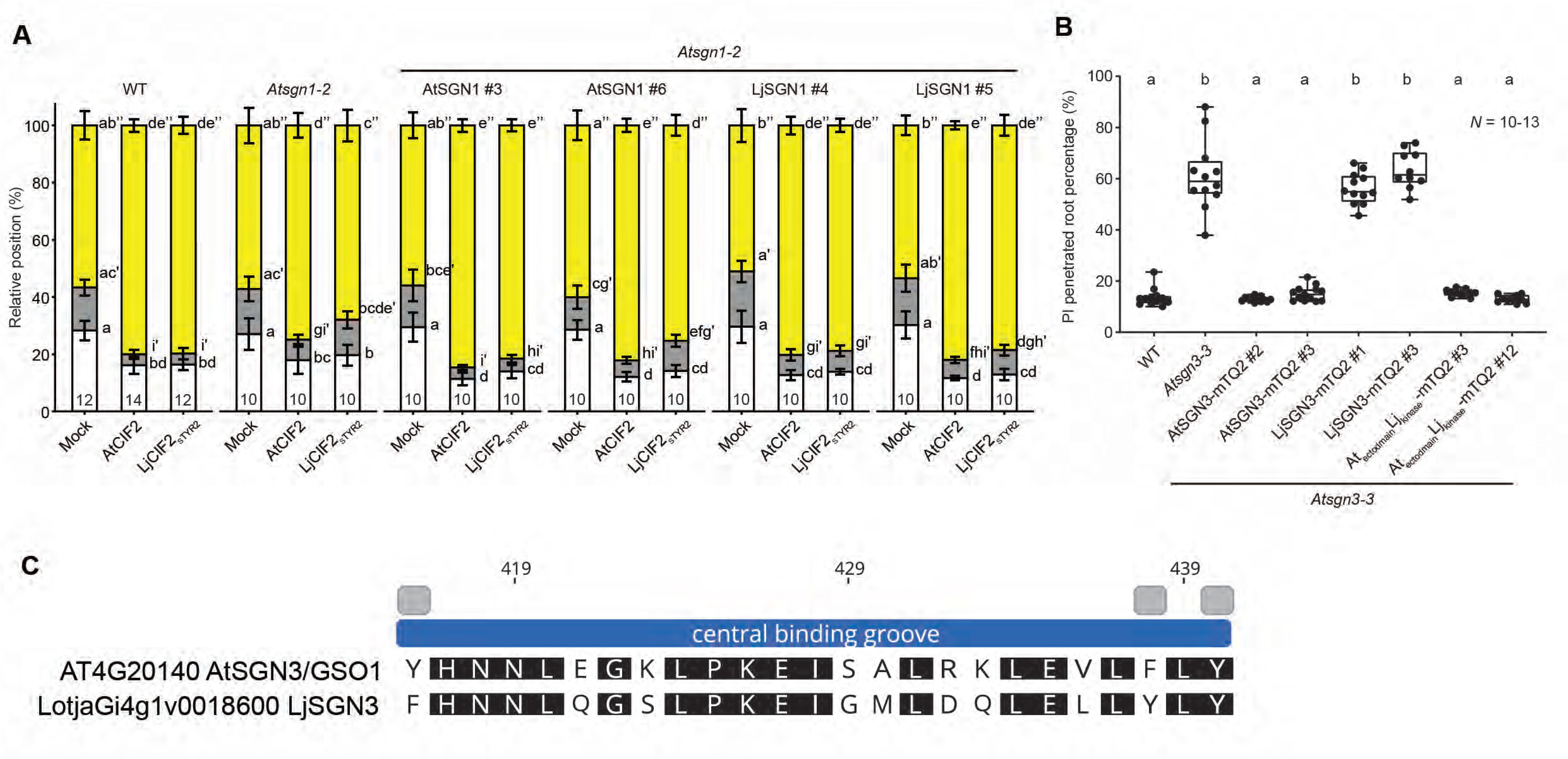
(**A**) Suberization pattern in 7-day-old roots of *Atsgn1-2* complementation lines treated with mock, AtCIF2, or LjCIF2_sTYR2_ for 48 h. (**B**) Quantification of propidium iodide (PI) penetration in 7-day-old roots of *Atsgn3-3* complementation lines. (**C**) Amino acid sequence alignment of AtSGN3 and LjSGN3 at the proposed central binding groove of ligand. Numbers correspond to the positions in full-length sequence of AtSGN3. Grey boxes represent validated amino acids in (*34*). Different letters in (**A**) and (**B**) depict the statistical difference in a one-way ANOVA analysis with Tukey’s test (*P* < 0.05).

**Supplementary Figure 5.**
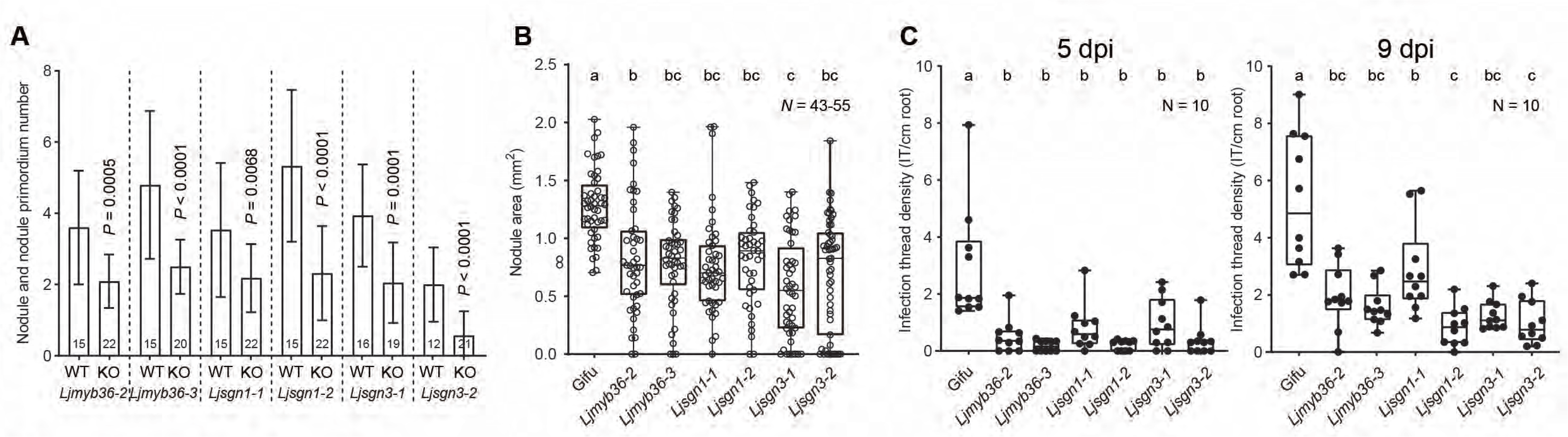
(**A**) Quantification of nodule and nodule primordium number at 21 days post-inoculation (dpi) after mass inoculation. *P* values indicate two-sided Student’s t test. (**B**) Quantification of nodule area at 21 dpi after mass inoculation. Each dot represents the biggest nodule formed by an individual plant. Note that there were cases that mutant roots did not form nodules. Combined results from two independent experiments. (**C**) Quantification of infection thread density at 5 dpi and 9 dpi after mass inoculation. Representative result from two independent experiments. Different letters in (**B**) and (**C**) depict the statistical difference in a one-way ANOVA analysis with Tukey’s test (*P* < 0.05), unless otherwise indicated.

**Supplementary Figure 6.**
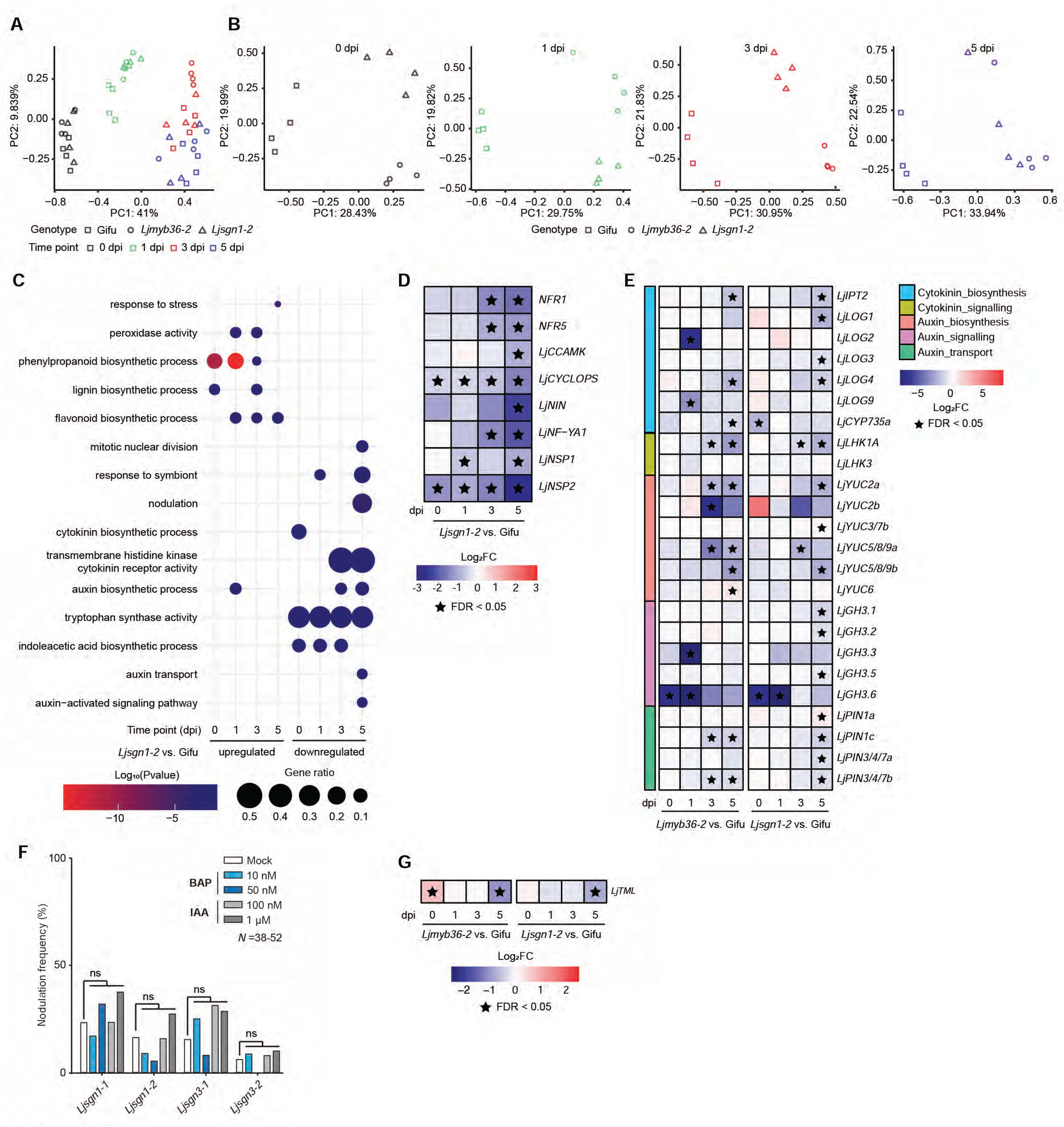
(**A**) Principal coordinate analysis (PCoA) plot of nodule primordium transcriptome of combined time points. (**B**) PCoA plot of nodule primordium transcriptome at individual time points. (**C**) Dotplot of Gene Ontology terms enriched in upregulated and downregulated differentially expressed genes of *Ljsgn1-2* nodule primordia at different time points after spot inoculation. (**D**) Heatmap of symbiosis related genes in *Ljsgn1-2* nodule primordia at different time points. (**E**) Heatmap of auxin and cytokinin related genes in *Ljmyb36-2* and *Ljsgn1-2* nodule primordia at different time points. (**F**) Quantification of occurrence of nodule formation at 14 days post-inoculation (dpi) after spot inoculation. Fisher’s exact test was performed between hormone treatment and mock within the same genotype (ns, not significant). Combined results from two independent experiments. (**G**) Heatmap of *TML* in *Ljmyb36-2* and *Ljsgn1-2* nodule primordia at different time points.

**Supplementary Figure 7.**
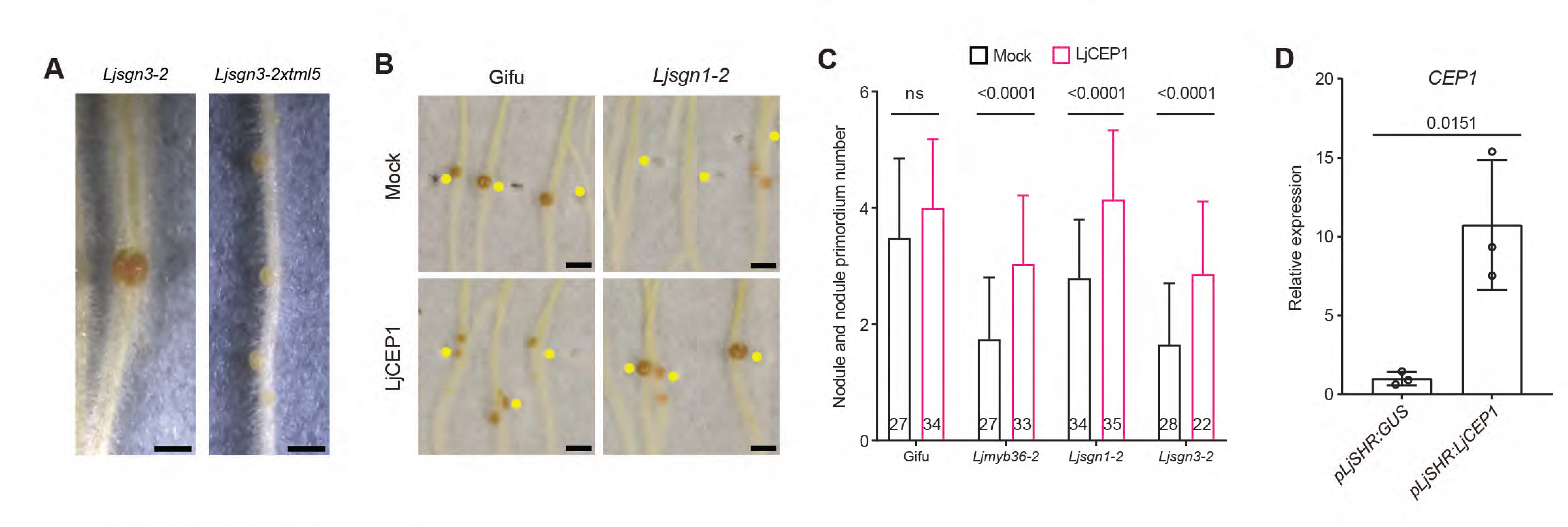
(**A**) Representative images of mass-inoculated roots with nodules at 28 days post-inoculation (dpi). (**B**) Representative images of mass-inoculated roots treated with mock (water) or 1 µM LjCEP1 at 21 dpi. Yellow dots indicate the position of primary root tip at 0 DAI. (**C**) Quantification of nodule and nodule primordium number on plants treated with mock (water) or 1 µM LjCEP1 on shoots at 21 dpi after mass inoculation. Numbers on the graph indicate the number of plants investigated. Representative results from two independent experiments. (**D**) Relative expression level of *LjCEP1* in *Ljmyb36-2* hairy roots expressing *pLjSHR:GUS* or *pLjSHR:LjCEP1*. Scale bar: 1 mm (**A**), 2 mm (**B**).

**Supplementary Figure 8.**
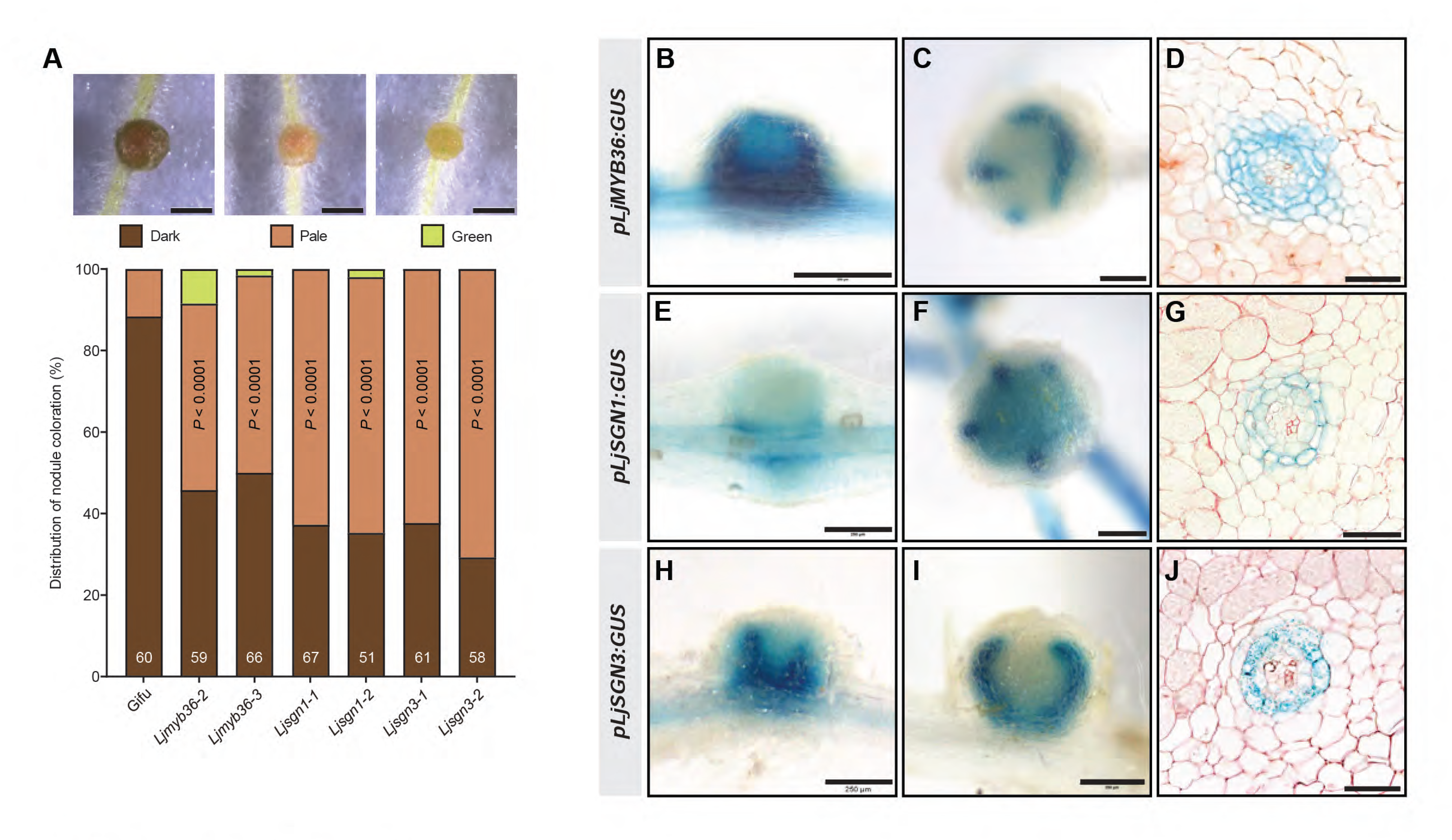
(**A**) Quantification of different nodule colorations at 21 days post-inoculation (dpi). Numbers on the graph indicate the number of plants investigated, combined results from three independent experiments. Note that only plants with mature nodules were included in the analysis. *P* values indicate Fisher’s exact test between Gifu and mutants regarding pale colored nodules. Transcriptional activity of *LjMYB36* in young nodule (12 dpi) (**B**), mature nodule (21 dpi) (**C**) and nodule vascular bundle (21 dpi) (**D**). Transcriptional activity of *LjSGN1* in young nodule (12 dpi) (**E**), mature nodule (21 dpi) (**F**) and nodule vascular bundle (21 dpi) (**G**). Transcriptional activity of *LjSGN3* in young nodule (12 dpi) (**H**), mature nodule (21 dpi) (**I**) and nodule vascular bundle (21 dpi) (**J**). Scale bar: 1 mm (**A**), 250 µm (**B,C,E,F,H,I**), 50 µm (**D,G,J**).

**Supplementary Figure 9.**
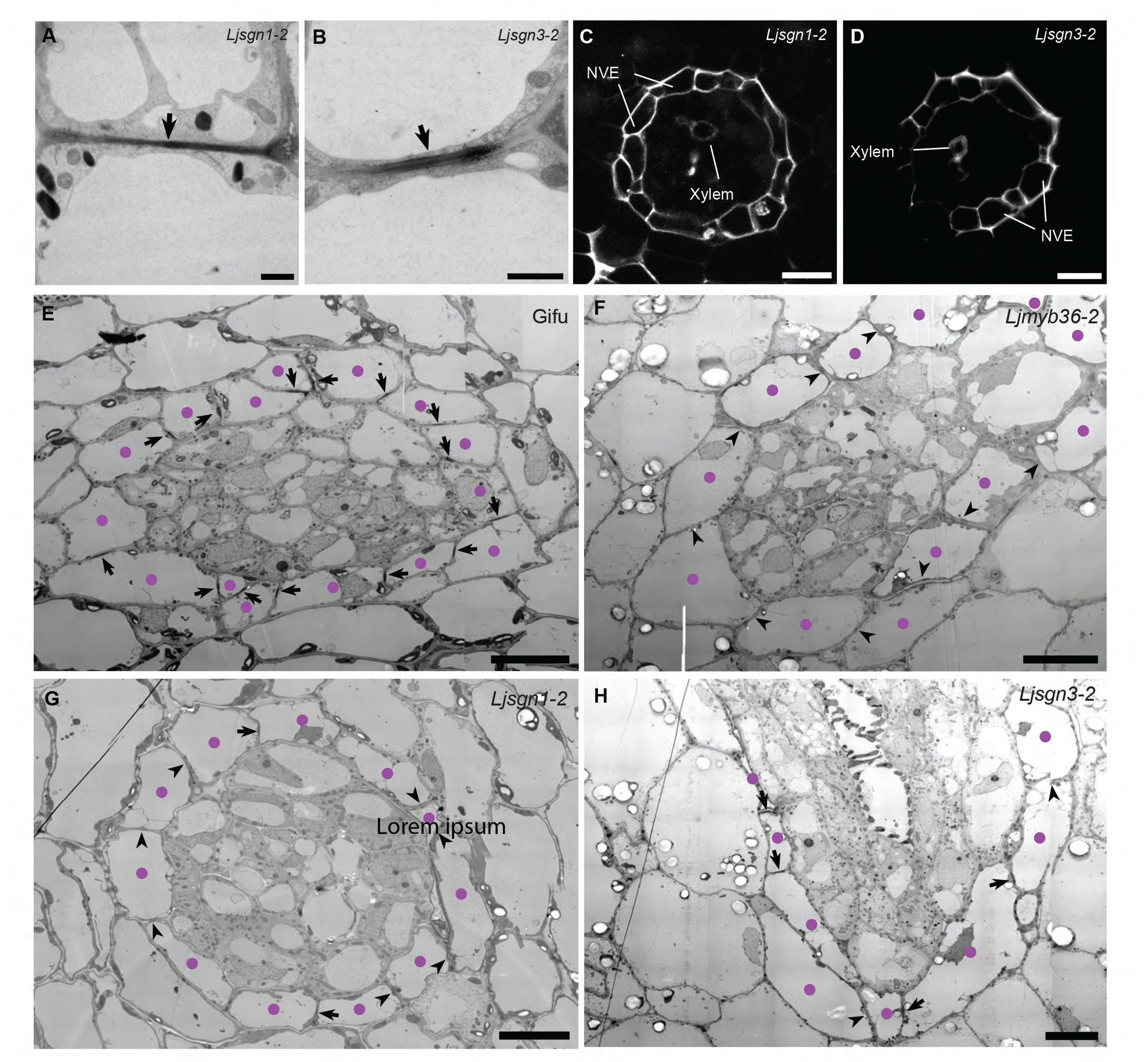
Transmission electron microscopy (TEM) images of *Ljsgn1-2* (**A**) and *Ljsgn3-2* (**B**) nodule vascular endodermis (NVE) at 21 days post-inoculation (dpi) (anticlinal wall between two endodermal cells). Electron-dense depositions highlighted by arrows represent Casparian strips (CSs) in NVE. Representative images of at least five nodules from individual plants. Nodule vascular endodermis stained with propidium iodide (PI) dye (white signal) in *Ljsgn1-2* (**C**) and *Ljsgn3-2* (**D**) at 21 dpi. Representative images of at least 10 nodules from individual plants. Overview TEM images of Gifu (**E**), *Ljmyb36-2* (**F**), *Ljsgn1-2* (**G**) and *Ljsgn3-2* (**H**) nodule vascular endodermis at 21 dpi. Purple dots indicate NVE. Arrows indicate CS, arrowheads indicate the absence of CS, when discernable. Scale bar: 1 µm (**A,B**), 20 µm (**C,D**), 10 µm (**E-H**).

**Supplementary Figure 10.**
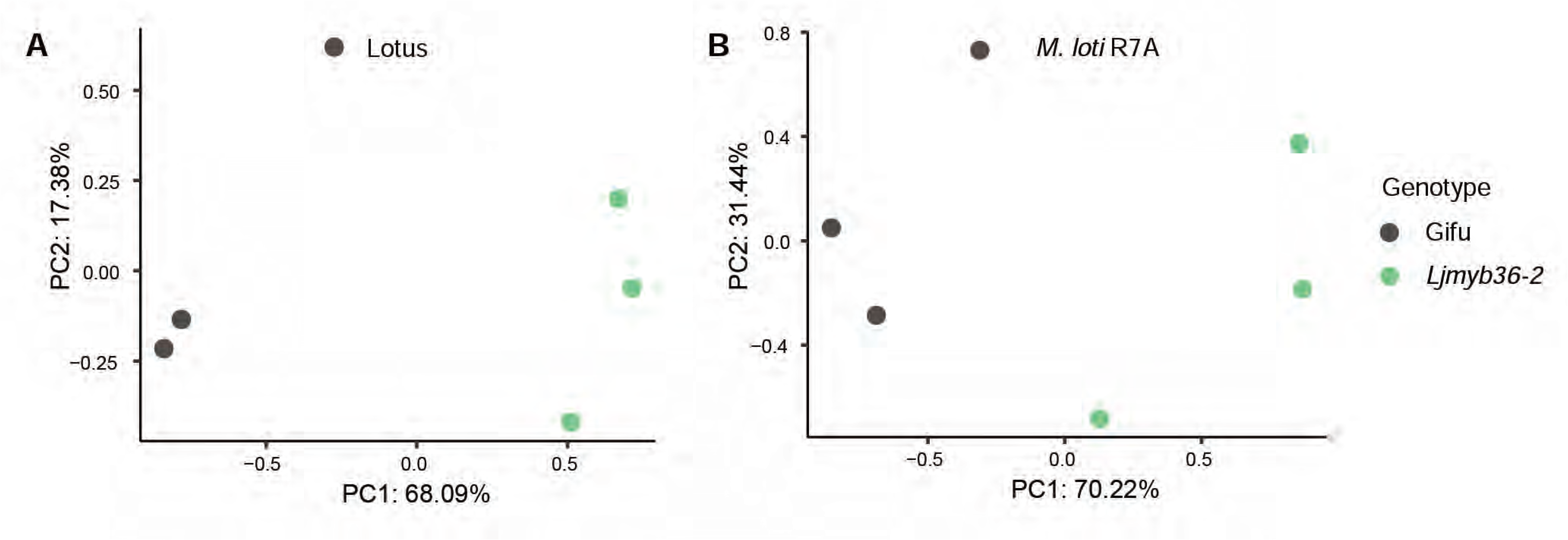
Principal coordinate analysis (PCoA) plot of Lotus (**A**) and *Mesorhizobium loti* R7A rhizobia (**B**) transcriptome.

**Supplementary Table 1.** Amino acid identity between *Lotus japonicus* and *Arabidopsis thaliana* orthologous.

**Supplementary Table 2.** Information of *LORE1* mutant lines generated in this study.

**Supplementary Table 3.** Primers used in this study.

**Supplementary Table 4.** Differentially expressed genes of RNA-seq analysis.

**Supplementary Table 5.** Acronyms with accession numbers used in this study.

## Notes

### Competing Interest Statement

The authors have declared no competing interest.

